# Genetic Analyses of Blood Cell Structure for Biological and Pharmacological Inference

**DOI:** 10.1101/2020.01.30.927483

**Authors:** Parsa Akbari, Dragana Vuckovic, Tao Jiang, Kousik Kundu, Roman Kreuzhuber, Erik L. Bao, Louisa Mayer, Janine H. Collins, Kate Downes, Michel Georges, Luigi Grassi, Jose A. Guerrero, Stephen Kaptoge, Julian C. Knight, Stuart Meacham, Jennifer Sambrook, Denis Seyres, Oliver Stegle, Jeffrey M. Verboon, Klaudia Walter, Nicholas A. Watkins, John Danesh, David J. Roberts, Emanuele Di Angelantonio, Vijay G. Sankaran, Mattia Frontini, Stephen Burgess, Taco Kuijpers, James E. Peters, Adam S. Butterworth, Willem H. Ouwehand, Nicole Soranzo, William J. Astle

**Affiliations:** British Heart Foundation Cardiovascular Epidemiology Unit, Department of Public Health and Primary Care, University of Cambridge, Strangeways Research Laboratory, Wort’s Causeway, Cambridge, CB1 8RN, UK; Department of Human Genetics, The Wellcome Sanger Institute, Wellcome Genome Campus, Hinxton, Cambridge, CB10 1HH, UK; Medical Research Council Biostatistics Unit, University of Cambridge, Cambridge Institute of Public Health, Cambridge Biomedical Campus, Forvie Site, Robinson Way, Cambridge, CB2 0SR, UK; The National Institute for Health Research Blood and Transplant Unit in Donor Health and Genomics, University of Cambridge, Strangeways Research Laboratory, Wort’s Causeway, Cambridge, CB1 8RN, UK; Department of Haematology, University of Cambridge, Cambridge Biomedical Campus, Long Road, Cambridge, CB2 0PT, UK; Division of Hematology/Oncology, Boston Children’s Hospital, Harvard Medical School, 1 Blackfan Circle, Boston, MA 02115, United States; Department of Pediatric Oncology, Dana-Farber Cancer Institute, Harvard Medical School, 450 Brookline Ave, Boston, MA 02215, United States; Broad Institute of MIT and Harvard, 415 Main St, Cambridge, MA 02142, United States; Harvard-MIT Health Sciences and Technology, Harvard Medical School, 77 Massachusetts Ave, Cambridge, MA 02139, United States; National Health Service Blood and Transplant, Cambridge Centre, Cambridge Biomedical Campus, Long Road, Cambridge, CB2 0PT, UK; Department of Haematology, Barts Health National Health Service Trust, Barts Health NHS Trust, London, E1 1BB, UK; Unit of Animal Genomics, GIGA-R & Faculty of Veterinary Medicine, University of Liège, B34, 1 avenue de l’Hôpital, B-4000, Liège, Belgium; National Institute for Health Research Cambridge BioResource, Box 229, Addenbrooke’s Hospital, Cambridge Biomedical Campus, Cambridge, CB2 0QQ, UK; Wellcome Centre for Human Genetics, University of Oxford, Roosevelt Drive, Oxford, OX3 7BN, UK; European Molecular Biology Laboratory, European Bioinformatics Institute, Wellcome Trust Genome Campus, Hinxton, Cambridge, CB10 1SD, UK; European Molecular Biology Laboratory, Genome Biology Unit, 69117 Heidelberg, Germany; Division of Computational Genomics and Systems Genetics, German Cancer Research Center (DKFZ), 69120, Heidelberg, Germany; British Heart Foundation Centre of Research Excellence, University of Cambridge, Cambridge, UK, Box 110, Addenbrooke’s Hospital, Cambridge Biomedical Campus, Cambridge, CB2 0QQ, UK; Health Data Research UK Cambridge, Wellcome Genome Campus and University of Cambridge, Cambridge, UK; Nuffield Division of Clinical Laboratory Sciences, Radcliffe Department of Medicine, University of Oxford, John Radcliffe Hospital, Headley Way, Headington, Oxford, OX3 9DU, UK; National Institute for Health Research Oxford Biomedical Research Centre—Haematology Theme, John Radcliffe Hospital, Headley Way, Headington, Oxford, OX3 9DU, UK; National Health Service Blood and Transplant, Oxford Centre, John Radcliffe Hospital, Headley Way, Headington, Oxford OX3 9BQ, UK; Department of Blood Cell Research, Sanquin Research and Landsteiner Laboratory, Sanquin, University of Amsterdam, Amsterdam, Netherlands; Department of Pediatric Immunology, Rheumatology and Infectious Disease, Emma Children’s Hospital, Amsterdam University Medical Center, Amsterdam, Netherlands; Department of Immunology and Inflammation, Imperial College London, Commonwealth Building, The Hammersmith Hospital, Du Cane Road, London, W12 0NN, UK

**Keywords:** hematology, blood cells, flow cytometry, genetics, intermediate traits, plasma proteome, *α*-granules, complex disease, drug target validation

## Abstract

Thousands of genetic associations with phenotypes of blood cells are known, but few are with phenotypes relevant to cell function. We performed GWAS of 63 flow-cytometry phenotypes, including measures of cell granularity, nucleic acid content, and reactivity, in 39,656 participants in the INTERVAL study, identifying 2,172 variant-trait associations. These include associations mediated by functional cellular structures such as secretory granules, implicated in vascular, thrombotic, inflammatory and neoplastic diseases. By integrating our results with epigenetic data and with signals from molecular abundance/disease GWAS, we infer the hematopoietic origins of population phenotypic variation and identify the transcription factor FOG2 as a regulator of platelet *α*-granularity. We show how flow cytometry genetics can suggest cell types mediating complex disease risk and suggest efficacious drug targets, presenting Daclizumab/Vedolizumab in autoimmune disease as positive controls. Finally, we add to existing evidence supporting IL7/IL7-R as drug targets for multiple sclerosis.

## INTRODUCTION

Blood cells have functions in a wide variety of physiological processes, most importantly in oxygen transport, hemostasis, and the immune system. There are three principal types of cells in the peripheral blood: red cells, which carry oxygen from the lungs for respiration; platelets, which initiate repair of damaged blood vessels to prevent bleeding; and white cells which clear pathogens through innate and acquired immune processes. These cells are suspended in plasma, an aqueous solution containing glucose, metabolites, hormones and proteins. Many of these proteins have a hematopoietic origin, because biomolecules from mature blood cells routinely diffuse into the plasma during functional processes such as cell activation or granule secretion.

Blood cell function cannot, at present, be measured using high throughput instruments. Consequently, genetic association studies of cell function have been limited to relatively small studies of platelet aggregation phenotypes, which have identified fewer than ten associated variants (Johnson *et al.* 2010). High powered genetic association studies of blood cell phenotypes have been performed, but these have concentrated on variables measured in classical complete blood counts (cCBCs) (Astle *et al.* 2016; Tajuddin *et al.* 2016; Chami *et al.* 2016; Eicher *et al.* 2016). cCBCs are standard clinical reports, which include standardised counts of the cells in the peripheral blood, measurements of the average volumes of platelets and red cells and measurements of blood hemoglobin concentration and localization. These variables contain a wealth of information about hematopoiesis and blood cell clearance mechanisms, but they were not selected to measure properties of the intracellular structures which participate in clinically relevant cell activation pathways. For example, pathologies of leukopoiesis that result in a lack of specific granules in neutrophils can cause immune dysfunction (Amulic *et al.* 2012), but such disorders are frequently accompanied by a normal neutrophil count and so are not detected by cCBCs.

We took a new approach to phenotyping biological variation endogenous to blood cells. Modern hematology analyzers (which generate cCBCs reports) contain flow cytometers which measure properties of blood cells by recording disturbances of incident laser light. The Sysmex XN-1000 instrument, for example, treats blood with membrane permeabilizing agents, allowing a nucleic acid staining dye to enter the cells. The fluorescence intensity of light emitted by the dye (termed ‘side fluorescence’ or SFL) measures the nucleic acid content and membrane permeability of the cell, while the intensities of diffracted light scattered sideways (SSC) and forward (FSC) by the cell measure intracellular organelle complexity and cell size respectively (**Figures 1A-E**) (Zimmermann *et al.* 2011; Linssen *et al.* 2008). When aggregated at the individual level (by median or distribution width), these data provide summary measures of cell structure related to cell function. For example, antigen-stimulated (reactive) lymphocytes can be detected and enumerated using SFL (Henriot *et al.* 2017) while the activation of neutrophils (NE) with physiological compounds such as formyl-methionyl-leucyl-phenylalanine (fMLP), and lipopolysaccharide (LPS) causes changes in NE-SSC (median neutrophil SSC) and NE-SFL (median neutrophil SFL) (Zimmermann *et al.* 2011; Linssen *et al.* 2008).

**Figure 1.**
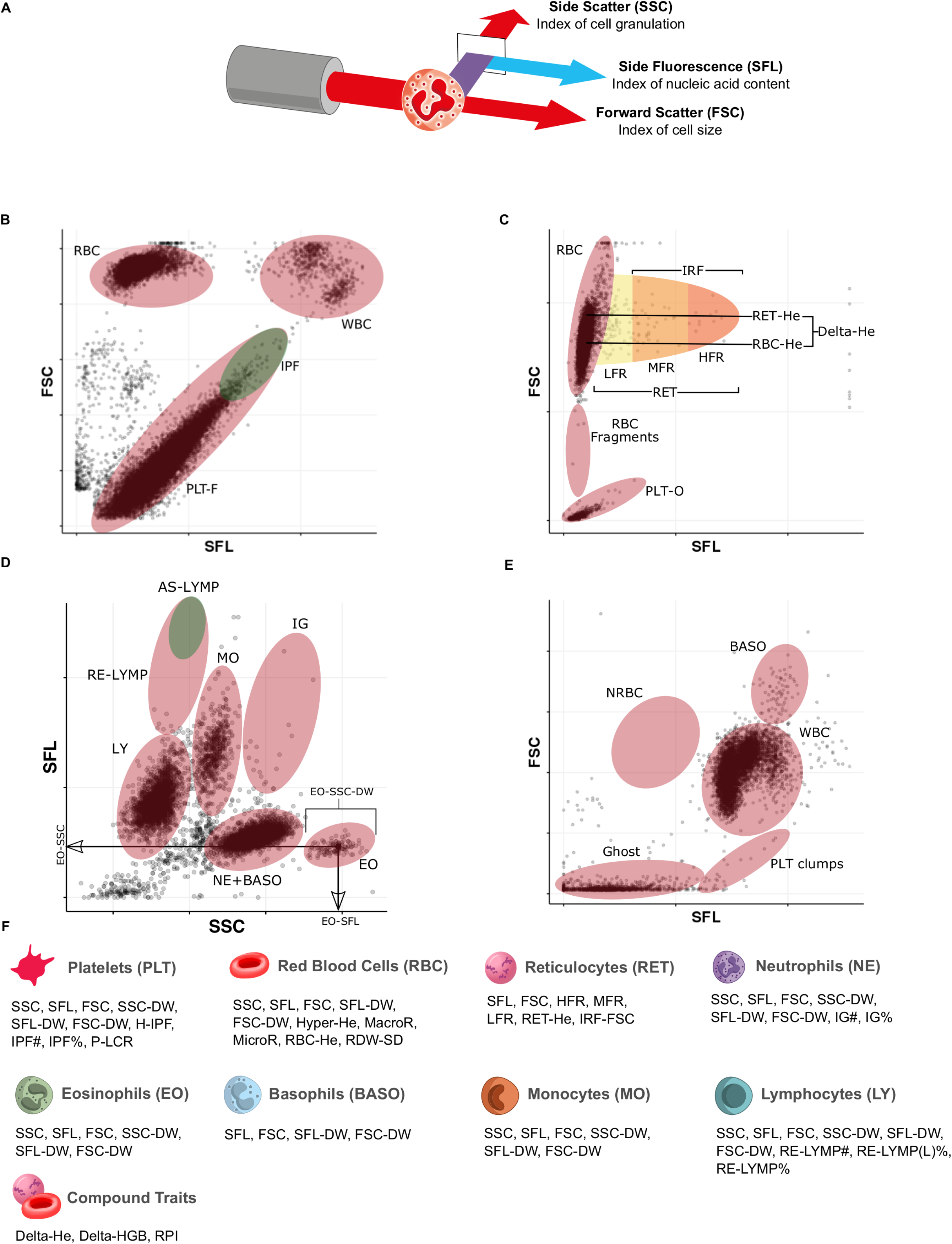
Flow Cytometry Traits Measured by the Sysmex XN-1000 Haematology Analyser (Adapted from (Sysmex 2014)) (A) Schematic of a granulocyte cell passing through the laser of the internal flow cytometer of the analyser. The instrument measures the intensities of incident light scattered sidewise (SSC, cell complexity/granularity) and forward (FSC, cell volume) by the cell and the intensity of the light which is absorbed and fluoresced at a new wavelength (SFL, cell nucleic acid content). (B)-(E) Cytometry scattergrams from an arbitrary participant: 2-dimensional projections of the cell level intensity data (SSC, SFL, FSC) measured in each of the four XN-1000 flow cytometry channels active for the INTERVAL study: PLT-F (platelet flow) channel (B), RET (reticulocyte) channel (C), WDF (white cell differential) channel (D), WNR (white cell and nucleated red cell) channel (E). Many of the traits correspond to medians or distribution widths (DWs) of clusters of cell level measurements (illustrated in panel (D) for three eosinophil traits) within the ellipses, which indicate the approximate regions occupied by cells of various types. Supplementary Table 1 contains a full description of the measurement procedure for each trait. (F) The 63 cytometry traits classified by the type of cells which they measure: platelets (PLT), mature red blood cells (RBC), reticulocytes (RET), neutrophils (NE), eosinophils (EO), basophils (BASO), monocytes (MO) and lymphocytes (LY). The three compound traits (Delta-HE, Delta-HGB, and RPI) depend on measurements of both mature red cells and reticulocytes.

Here, we report the first large-scale genome-wide association studies (GWAS) of flow cytometry measured non-classical CBC (ncCBC) traits (**Figure 1F**). We call these phenotypes ncCBC traits, because they were acquired using a hematology analyser (the XN-1000), but are not included in standard cCBC reports. By integrating the results of ncCBC, multi-omic and disease GWAS, we show how flow cytometry traits can be used to elucidate the secretory origins of proteins in the blood plasma, to study biological pathways mediating variation in disease risks and to suggest the cell types of action of drugs.

## RESULTS

### Hundreds of new genetic determinants of blood cell flow cytometry traits

We studied 63 ncCBC phenotypes, of which 11 summarise measurements of intracellular complexity/granularity (SSC), 16 summarise measurements of cell nucleic acid content/membrane lipid content (SFL) and 15 summarise measurements of cell morphology/volume (FSC) (**Table S1**). 20 of the traits were platelet phenotypes, 10 were red cell phenotypes and 33 were white cell phenotypes. The effective numbers of uncorrelated platelet, red cell and white cell traits (see STAR★Methods) were 5.2, 11.3 and 28.4 respectively (**Figures S1, S2**). Furthermore, the effective numbers of additional uncorrelated platelet, red cell and white cell traits, compared to those of the same cell type measured by cCBCs were 3.9, 2.7 and 24.7 respectively, suggesting that the white cell traits collectively measure a greater diversity of novel biology than the platelet and red cell traits (**Figures 2**, **S3, S4**). We performed univariable genetic association analyses (**Figure 1F**; **Table S1**), using genotypes imputed for 26.8 million variants in 39,656 European-ancestry blood donors in the INTERVAL study (Moore *et al.* 2014). Stepwise regression analysis identified 2,172 distinct (conditional *P*-value<8.31×10^−9^) variant-trait associations (**Table S2**). The application of a standard LD (*r*^2^>0.8) based greedy clumping algorithm, partitioned the 1,314 unique variants identified by the stepwise analysis (the *conditionally significant variants*) into 849 clusters (clumps), of which 231 contained a platelet associated variant, 211 a red cell associated variant and 432 a white cell associated variant (**Figures 2**, **S5**). 74 (32%) of the clumps containing a platelet trait associated variant contained no variant in high LD (*r*^2^>0.8) with any of the conditionally significant variants associated with a platelet trait studied by Astle *et al.* (2016), in the largest GWAS of cCBC phenotypes published to date. Analogously, 54 (26%) of the clumps containing a red cell associated variant and 289 (69%) of the clumps containing a white cell associated variant contained no variant in high LD with a variant associated with a phenotype of the respective cell types in the earlier study of cCBCs (**Figure S6**). The enrichment of novel associations with white cell traits may be partly explained by the fact that cCBCs include some phenotypes of platelets and red cells that capture cellular morphology (e.g. mean cell volumes), for which there are no white cell analogues. More interestingly, in comparison to platelets and red cells, white cells exhibit substantial biological complexity susceptible to measurement by flow cytometry. Specifically, white cells are nucleated and contain complicated intracellular organelles such as granules and vacuoles which differ according to cell type. Both red cells and platelets are anuclear, and only platelets contain granules.

**Figure 2.**
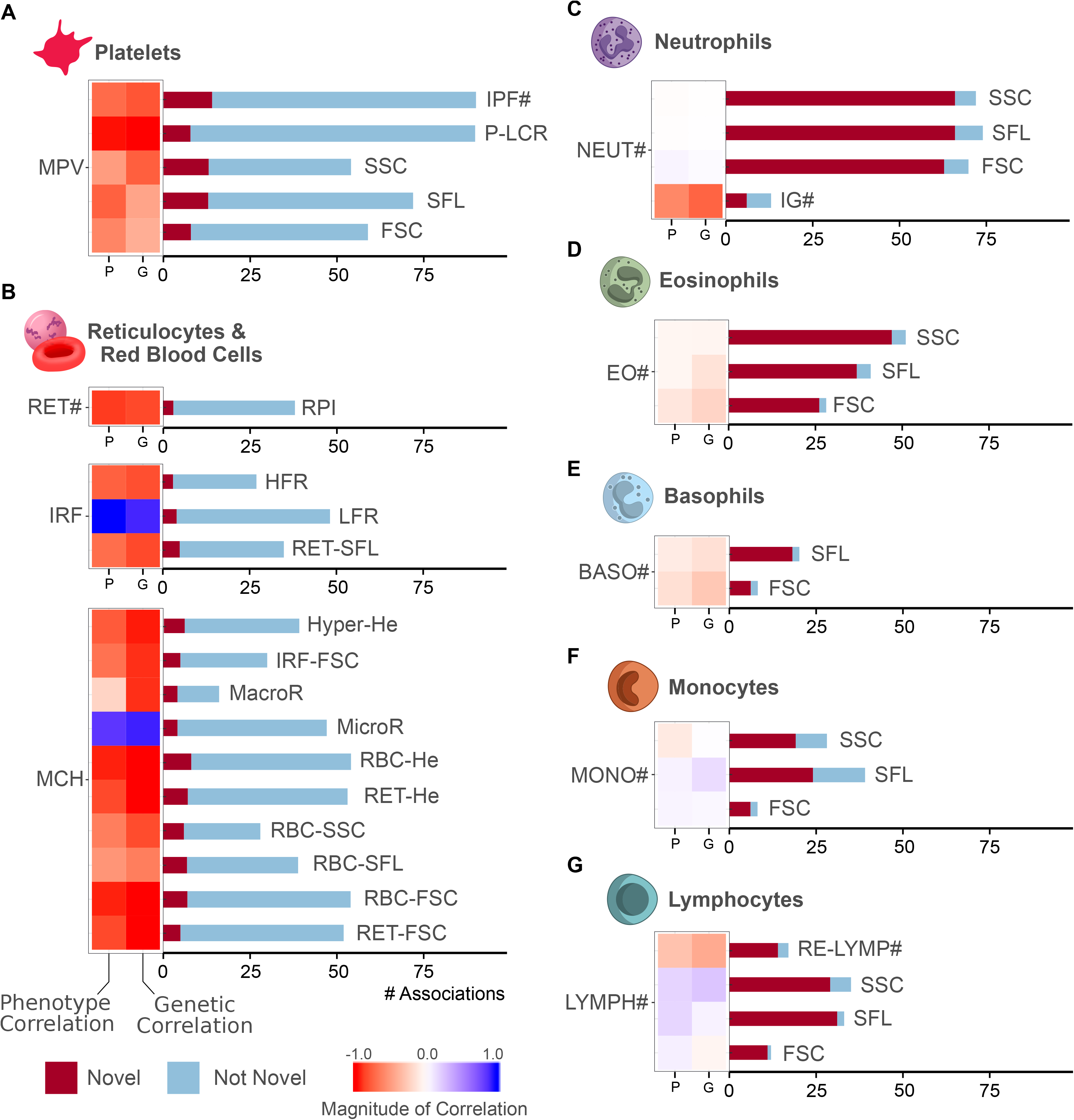
The Distribution and Novelty of Association Signals by Cell Type. (A)-(G) Each panel reports statistics for selected traits of the cell type indicated. The heat map on the left of each subplot shows the phenotypic (left) and genetic (right) correlation of the flow cytometry trait corresponding to each horizontal bar with the cCBC trait (those studied in Astle *et al.* (2016)) with which it has maximal absolute phenotypic correlation in the study sample. These cCBC traits have been used to group the ncCBC traits into separate subplots. The bar plot on the right of each subplot indicates the number of distinct (conditionally significant) associations for each cytometry trait and the number of those that share an LD clump with a variant reported to be associated with a blood trait of the same cell type by Astle *et al.* (‘Not Novel’). The white cell traits have lower genetic and trait correlation with their corresponding cCBC traits than the red cell and platelet traits, which is reflected in the distribution of new signals across traits.

### The effects of associated variants on transcription and cell structure

To identify possible molecular mediators of the association signals, we annotated each conditionally significant variant with the subset of its enveloping or proximal (within 5kb) genes for which the transcriptional consequence predicted by Variant Effect Predictor (VEP) was most severe in the Ensembl ranking (McLaren *et al.* 2016). We undertook an extensive search of the literature characterising these genes, which highlighted protein products with fundamental functions in the cell types corresponding to the associations, including thrombus formation for platelet traits (e.g. *VWF, SERINE2*), iron homeostasis for red cell traits (e.g. *HFE, TFRC*) and chemotaxis and adhesion for myeloid white cell traits (e.g. *P2RY2, SSH2*) (**Figure 3**; **Table S2**). A comparison of the genes assigned to the ncCBC associations with those assigned to cCBC associations by Astle *et al.* (using the same procedure) revealed that 25% of platelet, 25% of red blood cell, and 79% of white blood cell ncCBC gene assignments corresponded to novel associations with the corresponding cell type (red cells, platelets, neutrophils, eosinophils, basophils, monocytes, and lymphocytes). In total we identified 245 genes not reported by Astle *et al.*, 2016, including, for example, the eosinophil phenotype associated genes *CTNS* and *IFI30* (which respectively code for cystinosin, a lysosomal cystine transporter and gamma-interferon-inducible lysosomal thiol reductase) (West and Cresswell 2013; Guo *et al.* 2018).

**Figure 3.**
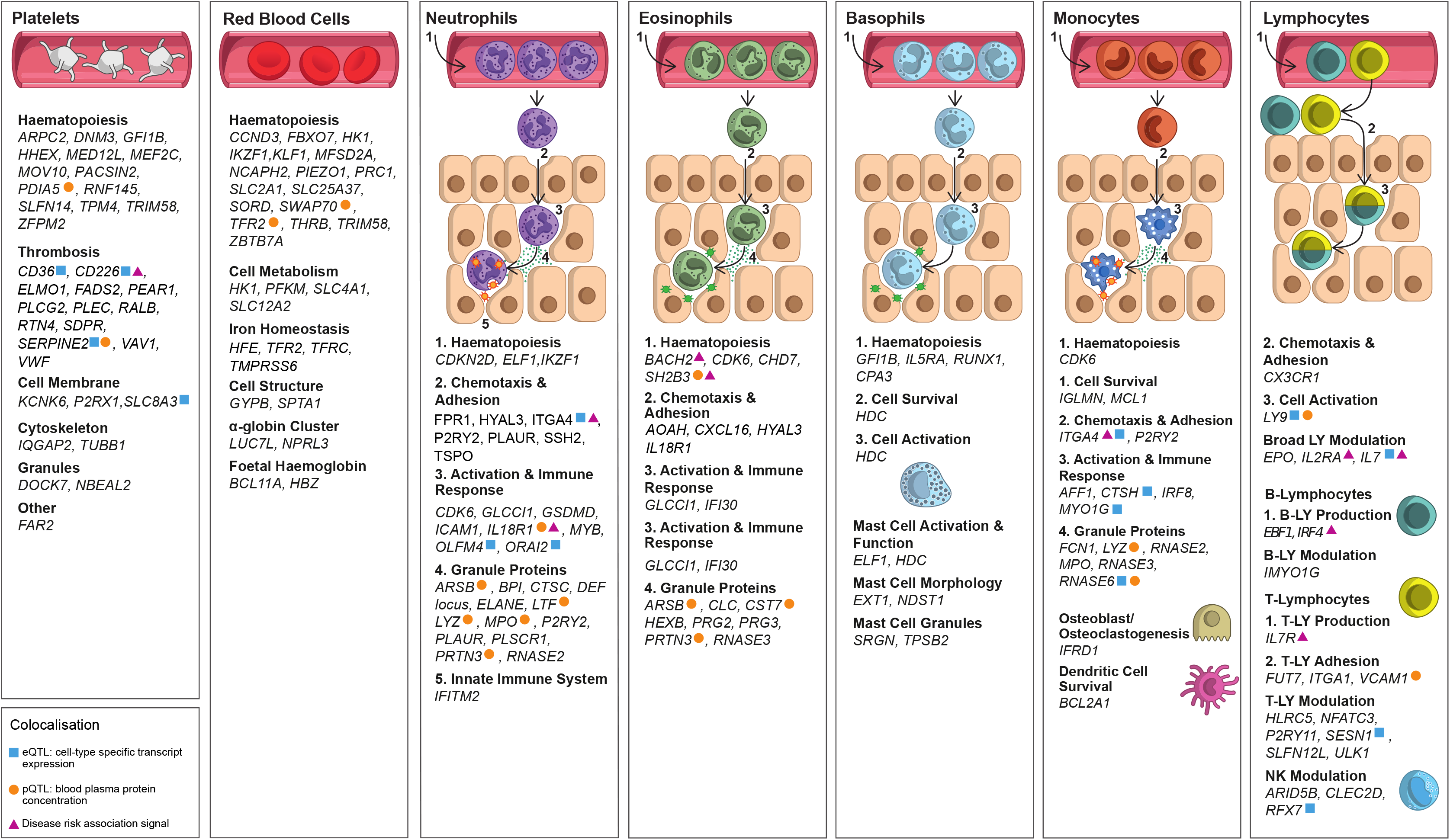
Summary of Biological Functions of VEP Assigned Genes Identified by a Survey of the Literature. Each panel contains a list of genes, assigned by VEP or by eQTL/pQTL colocalization, to genetic associations with traits corresponding to the given cell type, for which a literature search identified evidence of known function. Each list is stratified into functional categories relevant to the cell type. Table S2 contains a complete list of conditionally significant associated variants, their VEP annotated genes, and relevant references to literature. The coloured symbolic annotations indicate genes assigned to variants which colocalise with eQTL (blue square), pQTL (orange circle), or disease GWAS associations (purple triangle). An eQTL or pQTL colocalization may be with a cis signal for a nearby gene, which may differ from the gene assigned by VEP.

To assess the usefulness of the VEP annotation, we computed posterior probabilities (PPs) of colocalization between the ncCBC trait association signals and gene expression quantitative trait loci (eQTL) identified in the corresponding cell types. We aggregated imputed genotypes with microarray measurements of gene expression in platelets, neutrophils, monocytes, and CD19^+^ B, CD4^+^ T and CD8^+^ T lymphocytes from five independent studies containing 300-1,490 participants depending on cell type. We searched for variants associated with gene expression in gene specific windows (limited to the gene body and a 1MB extension from each gene boundary) (Kreuzhuber 2018). After applying stringent analysis criteria (see STAR★Methods) we identified 134 ncCBC variant-trait associations exhibiting strong evidence (PP > 80%) of colocalization with at least one eQTL, corresponding to a total of 65 genes (**Table S3**). VEP assigned a concordant gene to 79 (81%) of the 97 eQTL colocalizing variant-trait associations unique to a particular cell type. The colocalizing genes include several with well understood biological functions (e.g. *ARHGEF3, CD226, CD36*) (**Figure 3**), some consistent with a computationally inferred function imputed from protein-protein interaction networks (e.g. *HABP4*) and others for which knowledge of function is limited (e.g. *SEC14L5, ZMAT3*).

We used the Gene Ontology database (The Gene Ontology Consortium 2019) and data from mass spectrometric profiling of neutrophil granules (Rorvig *et al.,* 2013) to identify the subcellular localisation of the products of many of the ncCBC trait associated genes (**Table S4**). This revealed that proteins mediating the genetic associations reside in a wide variety of organelles including the nucleus, the mitochondria, the endoplasmic reticulum, the Golgi apparatus, the centrosome, vesicles, lysosomes, vacuoles and cytoplasmic granules. Cellular granules are organelles, which are prominent in neutrophils, eosinophils, basophils, and present in monocytes where they play critical roles in innate immune responses by releasing antimicrobial proteins (Borregaard, Sørensen, and Theilgaard-Mönch 2007; Acharya and Ackerman 2014; Becknell *et al.* 2015; MacGlashan 2013). We identified genes that code for a number of white cell granule proteins not previously found by GWAS of granulocyte traits, including Arylsulfatase B, Bactericidal permeability-increasing protein, RNase 3 and RNase K6. These proteins have well understood roles in immunity (Borregaard, Sørensen, and Theilgaard-Mönch 2007), illustrating a limitation of cCBC GWAS as a tool to study the functional properties of blood cells. More generally, we identified 48 associations with NE-SSC (neutrophil granularity), 39 associations with EO-SSC (eosinophil granularity), and 16 associations with MO-SSC (monocyte granularity) that did not correspond to any of the genetic associations with cCBC traits reported by Astle *et al.* (**Table S2**). These include associations near genes the functions of which in blood cells are well understood, including roles in transcription and translation (*BCL6, E2F2, EVI5*), in granule formation and retention (*VAMP3*, *VPS26A*), as granule cargo (*ARSB, CTSB, DEFA, PRTN3, PRG2/3, RNASE2/3*), and in post-translational modifications (*B3GNT2*, *CHST11*, *DSTYK*). Other associations highlight genes for which knowledge of hematological function is still developing (e.g. *PTBP1*, Polypyrimidine tract binding protein 1, which may modulate pre-mRNA processing (Attig *et al.* 2018) and is associated with NE-SSC; *TRPS1*, transcriptional repressor GATA binding 1, which functions as a transcriptional repressor, is implicated in human cancers (Wang *et al.* 2018) and is associated with EO-SSC; *VMP1*, vacuole membrane protein 1, which is required for organelle biogenesis, protein secretion, and autophagy (Calvo-Garrido *et al.* 2008; Zhao *et al.* 2017) and which is associated with MO-SSC), confirming that our approach has uncovered biologically interesting candidate genes and molecular pathways modulating granule formation and cellular complexity.

### The biogenesis of cellular structures

Many of the intracellular structures generating variation in flow cytometry traits have their origins in the immature precursors of peripheral blood cells. Granule formation for example, is a cell-type specific process known to occur at particular stages of cellular differentiation; the granules of platelets and granulocytes begin to form respectively in megakaryoblasts and myeloblasts. Consequently, the absence of a colocalising mature blood cell eQTL for a cytometry trait association may reflect the fact that the associated genetic variant exerts its effects after lineage commitment, but before terminal differentiation. To test this hypothesis, we applied FINEMAP v1.3.1 (Benner *et al.* 2016) to identify credible sets of variants causally associated with the ncCBC traits and assessed the enrichment of those variant sets in the nucleosome depleted regions (ATAC-seq) of nine types of progenitor cell (localized in the bone marrow) and nine types of mature cell (generally localized in the peripheral blood) (Ulirsch *et al.* 2019). We observed patterns of enrichment in the open chromatin regions of the progenitor cell types ancestral to neutrophils, eosinophils, monocytes, and lymphocytes (**Figure S7A-D**), which suggest the developmental stage (**Figure S7F**) at which the cell characteristics corresponding to particular traits (SSC, SFL, FSC) develop. For instance, neutrophils, which are the primary anti-microbial cell type, contain three classes of cytotoxic granules - azurophilic, specific and gelatinase - which are formed sequentially at distinct stages of differentiation (Grassi *et al.* 2018). The relative enrichments of genetic variants associated with NE-SSC (neutrophil granularity) in the nucleosome depleted regions of the hematopoietic stem cell (HSC) and the four types of myeloid progenitor cell (CMP, GMP-A, GMP-B, GMP-C) are consistent with a progressive increase in the accessibility of enhancers regulating granule formation during myeloid differentiation and point to an origin of these granules in lineage-committed myeloid progenitors (**Figure S7A-F**). In monocytes, we observed an enrichment of MO-SFL (monocyte nucleic acid content) associated variants in the nucleosome depleted regions of monocyte progenitor cells (GMP) consistent with the phase of proliferation and cell division, and an enrichment of MO-SSC (granularity) in the nucleosome depleted regions of peripheral blood monocytes indicating that granularity is regulated in the ultimate stages of monocyte differentiation before egress from the bone marrow.

### The cellular origins of plasma proteins

We hypothesized that some genetic variants associated with cytometric traits, in-particular genetic variants associated with side scatter traits, which can capture the abundance of secretory granules in cells, also influence the concentration of proteins in the blood plasma. To explore this, we turned to (Sun *et al.* 2018), a multi-GWAS study which identified 1,927 associations (protein quantitative trait loci, pQTL, **Table S4**) with the plasma concentrations of 1,478 proteins. For 943 of these proteins, there is strong evidence that transcripts are expressed (log scaled fragments per kilobase of transcript per million mapped reads (log_2_FPKM) >1.0) in at least one of the blood cell types surveyed by the ncCBC traits (megakaryocytes (MKs) and erythroblasts, the respective progenitors of platelets and red cells, and neutrophils, eosinophils, basophils, monocytes, CD4^+^ T cells, CD8^+^ T cells, naive B cells). We performed colocalization analysis between the pQTL and our cytometry trait-associated variants and identified 61 and 1,021 colocalizations (PP>80%) in cis and trans respectively (**Figure 3**, **Table S4**). 223 plasma proteins had associations colocalizing with just one cell-type suggesting that the hematopoietically derived fraction of these proteins in the plasma originates predominantly from a single blood cell type (e.g. *VEGFA*, which encodes vascular endothelial growth factor A, is associated with eight different platelet traits, while *RNASE6*, which encodes RNase K6, is associated solely with monocyte side fluorescence). Notably, many signals assigned by VEP to genes encoding granule proteins also colocalized with pQTL signals in blood plasma (**Figure 3**). Examples include *ARSB* (PP=99%)*, CST7* (PP=98%)*, CTSA* (PP=99%)*, LY9* (PP=98%)*, MPO* (PP=100%)*, PRTN3* (PP=99.9%) and *RNASE6* (PP=99%) (**Table S5**). The *PRTN3* colocalization illustrates how the genetic associations with cellular and molecular phenotypes can be combined to elucidate disease pathogenesis. rs138303849-C, a variant upstream of *PRTN3* (encoding proteinase-3, PR3), is known to be associated with an increased risk of a vasculitis characterised by autoantibodies against PR3 (Merkel *et al.* 2017). This variant is an eQTL in whole blood (GTEx Consortium 2015) and we recently showed that the risk allele is associated with increased PR3 concentration in plasma (Sun *et al.* 2018). The main source of PR3 in plasma is the specific granule, one of three classes of granule found in neutrophils. In the present analysis, we observed that the risk allele is associated with increased neutrophil SSC (granularity), SFL (nucleic acid), FSC (volume), and decreased NE-SSC-DW (SSC distribution width); the associations all colocalise with the cis-pQTL for higher plasma PR3 levels (PP=99.9%). This shows that the vasculitis risk allele not only increases *PRTN3* transcription and PR3 plasma protein levels, but also changes the structural and functional properties of neutrophils.

### Evidence that *ZFPM2* is a regulator of platelet *α*-granularity

Platelet activation is important in thrombus formation, wound healing, inflammation and the chemotaxis and activation of myeloid white cells. Critical to these biological processes are coagulation proteins, growth factors, proteases, chemokines and other signalling peptides that diffuse into the blood plasma when *α*-granules are released by activated platelets. Our GWAS of PLT-SSC (platelet granularity) identified an association in *ZFPM2* (which encodes the transcription factor Friend of GATA-2 or FOG2) (**Figure 4A**), colocalizing with trans pQTLs for 24 plasma proteins (*ANGPT1\**, *APLP2\**, *BDNF\**, *CCL17*, *COCH*, *CPXM1*, *CTSA\**, *DKK1*, *DKK4*, *DNAJB11*, *EDAR*, *ERP44*, *LGALS7*, *NSG2*, *PDGFA\**, *PDGFB\**, *PDGFD\**, *PPBP\**, *SERPINE1\**, *SIRT5*, *SPARC\**, *SYT11*, *UGT2A1*, *VEGFA\**) of which 11 (starred) are platelet *α*-granule localized proteins (**Table S5**). RNA-seq of 9 differentiated but nucleated blood cell types (Grassi *et al.* 2019) showed that transcripts of *ZFPM2* are substantially expressed solely in MKs (**Figure 4B**). Complementary data, from a study of the entire hematopoietic system, show that *ZFPM2* is specifically expressed in the MK lineage (**Figure 4C**). Stepwise multiple regression analysis of the platelet granularity association (PLT-SSC) suggested a single conditionally significant association signal in an 82kb interval of low recombination. Fine-mapping of the locus identified an intronic SNP (rs6993770) as the most probable causal variant (PP=95%). rs6993770 is located in a region of open chromatin (ATAC-seq) in MKs, which contains histone modifications indicative of enhancers (H3K4 tri-methylation, H3K27 acetylation; **Figure 4D, 4E**). The variant is 25bp upstream of a GATA motif on the negative strand, and 34bp upstream from the palindromic E-box binding motif CAGCTG. The juxtaposition of these motifs is characteristic of a hematopoietic co-binding site for GATA1 and TAL1, two of the three key megakaryocyte lineage determining transcription factors (Moreau *et al.* 2016; Han *et al.* 2016; Kassouf *et al.* 2010). None of the seven variants in high LD (*r*^2^ > 0.9) with rs6993770 were located in regions for which epigenetic data supported causality in MKs (**Figure 4E**).

**Figure 4.**
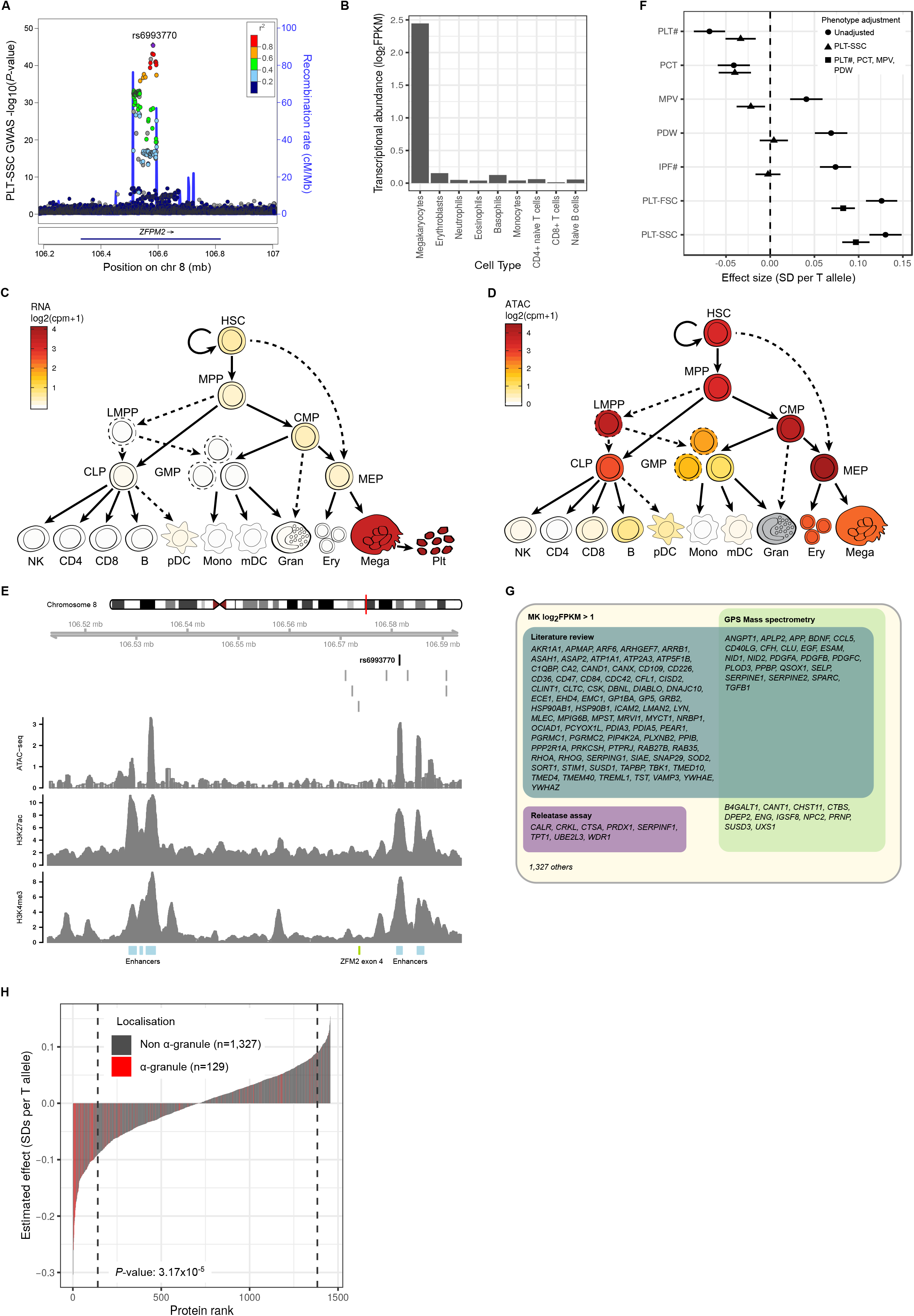
*ZFPM2* is a Regulator of Platelet *α*-granularity. **A.** LocusZoom plot for the *ZFPM2* locus (Pruim *et al.* 2010). Each dot corresponds to a variant in the locus. The *x*-axis shows physical position on chromosome 8 according to GrCh37 and the *y*-axis the −log_10_(*P*-value) from a univariable test for association between each variant’s imputed alternative allele count and PLT-SSC. The colour of the dot corresponds to the strength of correlation (*r*^2^) in the study sample with rs6993770, the sentinel variant. Conditional analysis identified a single association signal in the 82kb interval of low recombination containing rs6993770. **B.** Abundance of *ZFPM2* transcripts (log_2_FPKM) in megakaryocytes, erythroblasts, neutrophils, eosinophils, basophils, monocytes, CD4+ naive, CD8+, and naive T cells, showing that of these cell types *ZFPM2* transcription is limited to megakaryocyte cells (Grassi *et al.* 2019). **C.***ZFPM2* transcript expression is higher in platelets, megakaryocyte and relevant precursor cells MEP (megakaryocyte–erythroid progenitor cell), CMP (common myeloid progenitor), MPP (multipotent progenitor), and HSC (hematopoietic stem cell) compared to other blood cells and blood cell precursors. **D.** The *ZFPM2* locus remains accessible as assessed by ATAC-seq in a range of blood cell precursors, in particular platelet and megakaryocyte precursor cells MEP, CMP, MPP, and HSC. **E.** Epigenetic activity in MKs across the 82kb recombination interval containing the association signal. The *x*-axis shows physical position on chromosome 8. The location of rs6993770 is indicated by the dark vertical line, the nearby light vertical lines indicate the locations of seven variants in high LD (*r*^2^ > 0.9) with rs6993770. The *y*-axis of each panel measures sequencing read depth from an epigenetic assay. From top to bottom the panels correspond to ATAC-seq (open chromatin), H3K27ac (a mark of active enhancers) and H3K4me3 (a mark of accessibility to transcription factors). The blue rectangles at the bottom of the figure indicate the enhancer regions inferred from a set of six histone modifications (H3K4me1, H3K4me3, H3K9me3, H3K27ac, H3K27me3 and H3K36me3) in MKs using the IDEAS chromatin segmentation algorithm (Zhang *et al.* 2016; Petersen *et al.* 2017). The green rectangle indicates the position of exon 4 of *ZFPM2.* **F.** Forest plot showing the per allele effect of rs6993770-T on the means of the inverse rank normalized distributions of the platelet traits PLT# (platelet count), PCT (plateletcrit), MPV (mean platelet volume), PDW (platelet distribution width), IPF# (immature platelet fraction count), PLT-FSC (platelet volume), and PLT-SSC (platelet granularity) measured in the INTERVAL study. Circular symbols indicate estimates of the marginal effect, triangular points indicate estimates of the effect adjusted for PLT-SSC, square symbols indicate estimates of the effect adjusted for PLT#, PCT, MPV, and PDW. Horizontal lines correspond to 95% confidence intervals. The effect of rs6993770-T on PLT-SSC and PLT-FSC does not appear to be substantially mediated through the cCBC parameters. **G.** Venn diagram cross classifying the 1,456 genes that are expressed in MKs (transcript log_2_FPKM>1) and code for plasma proteins significantly (*P-*value < 10^−3^) associated with rs6993770-T. The classifying categories indicate a) implicated as an *α*-granule protein coding gene by a literature review b) a protein coded for by the gene was undetected by mass spectrometry of gray platelet syndrome patients’ platelets (which lack *α*-granules) but detected in healthy volunteers and c) protein expelled from activated platelets (platelet releasate). **H.** The estimated per allele effect of rs6993770-T on the mean concentration of 1,456 plasma proteins with coding genes transcribed in MK cells (log_2_FPKM>1). The *y*-axis measures the per allele effect size and the *x*-axis its rank. Bars corresponding to proteins located in platelet *α*-granules are colored red. Proteins with ranks in the tails bounded by the dashed lines exhibit significant evidence for an association with rs6993770-T at a relaxed critical threshold (*P-*value < 10^−3^). *α*-granule proteins are significantly (embedded *P*-value) enriched in the negative compared to the positive tail.

There are well known associations between rs6993770 and the four classical platelet traits: platelet count (PLT), volume (MPV), crit (PCT) and volume distribution width (PDW). (Gieger *et al.* 2011; Astle *et al.* 2016). We identified new associations with the ncCBC traits IPF#, PLT-SSC, and PLT-FSC (**Table S2, Figure 4F**). The estimated effect of rs6993770 on PLT-SSC was not significantly attenuated by a multivariable linear adjustment for the four classical traits, suggesting that the association is not substantially mediated through these traits. Consequently, we hypothesized that *ZFPM2* has a role in *α*-granule control. To examine this, we performed a broad analysis of the plasma concentrations of 1,456 of the proteins studied by (Sun *et al.* 2018) for which there was evidence for expression in MKs (RNA-seq log_2_FPKM>1, **Figure 4G**) and identified 215 proteins associated with rs6993770 at a relaxed univariable significance threshold (*P*-value<10^−3^). 44 of these 215 proteins are localized in *α*-granules and the estimated effect of rs6993770-T on the plasma concentration of 91% (40) of these 44 was negative. In contrast rs6993770-T had a negative estimated effect on the plasma concentration of 59% (101) of the 171 proteins not localised in *α*-granules, showing that the direction of effect of rs6993770-T on the plasma concentration of a protein differs significantly according to its *α*-granule localisation (Fisher’s exact test *P*-value=3.17×10^−5^) (**Figure 4H**). To assess whether this differential effect might be explained by a systematic difference in the expression level of genes according to the *α*-granule localisation of the corresponding proteins, we performed a linear regression of the estimated effect sizes of rs6993770-T on plasma concentration against a dummy variable indicating *α*-granule localisation, restricting to the 1,456 proteins expressed in MKs (log_2_FPKM>1) and adjusting for the abundance of the mRNA transcripts for the corresponding genes in MKs. We estimated the additive allelic effect of rs6993770-T on plasma proteins to be 0.369 phenotypic standard deviations lower on average (*P*-value 8.5×10^−10^) in proteins localized to *α*-granules compared to other proteins (**Figure 4H**; **Figure S8**). This shows that rs6993770 differentially modifies the plasma concentration of *α*-granule proteins, possibly by altering *ZFPM2* transcript levels in the cellular progenitors of platelets.

Interestingly, Klarin *et al.* (2017) recently reported an association between the T allele of rs6993770 and decreased risk of venous thromboembolism (VTE). They postulated that the underlying mechanism is a *ZFPM2* mediated decrease in the plasma concentration of the principal inhibitor of plasminogen activator PAI-1, which is encoded by *SERPINE1* (Klarin *et al.* 2019). The notion that lower levels of plasma PAI-1 cause a reduced risk of VTE is biologically plausible. However, we have shown that rs6993770 is pleiotropic, modifying the process of platelet formation and platelet granule content and the concentrations of many platelet derived proteins in the blood plasma (**Figure 4F**), highlighting the importance of broader multi-trait and multi-omic analysis, for etiological inference and the characterization of disease risk association signals.

### Elucidating disease etiology and inferring drug targets

We hypothesized that genetic associations with ncCBC traits could improve our understanding of etiological cell types and molecular mechanisms more generally. We chose to focus on immune, inflammatory and cardiovascular diseases, in which blood cells are known to play a causal role. We retrieved publicly available GWAS summary statistics for 26 diseases (Kundu *et al.* 2020) and assessed the evidence for colocalization between genetic associations with ncCBC traits and disease risks (**Table 1**) (Pickrell *et al.* 2016). We found strong evidence (PP>80%) for colocalization between 73 of the variant-ncCBC trait associations (in 29 LD clumps) and at least one disease association (**Table S6**).

**Table 1.**
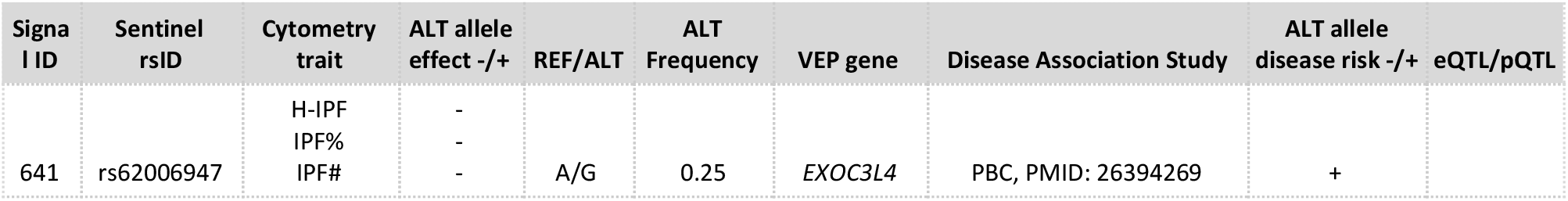
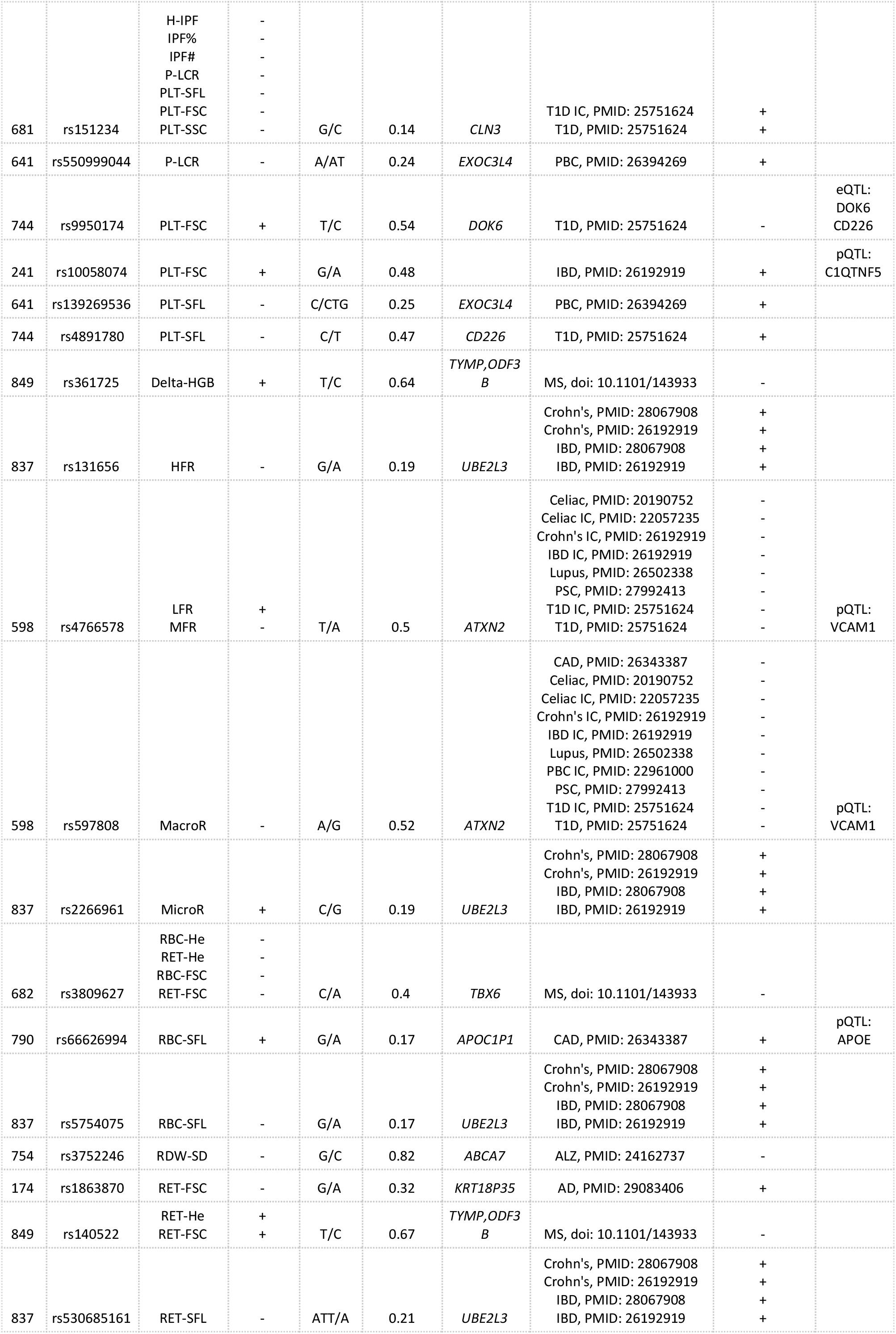
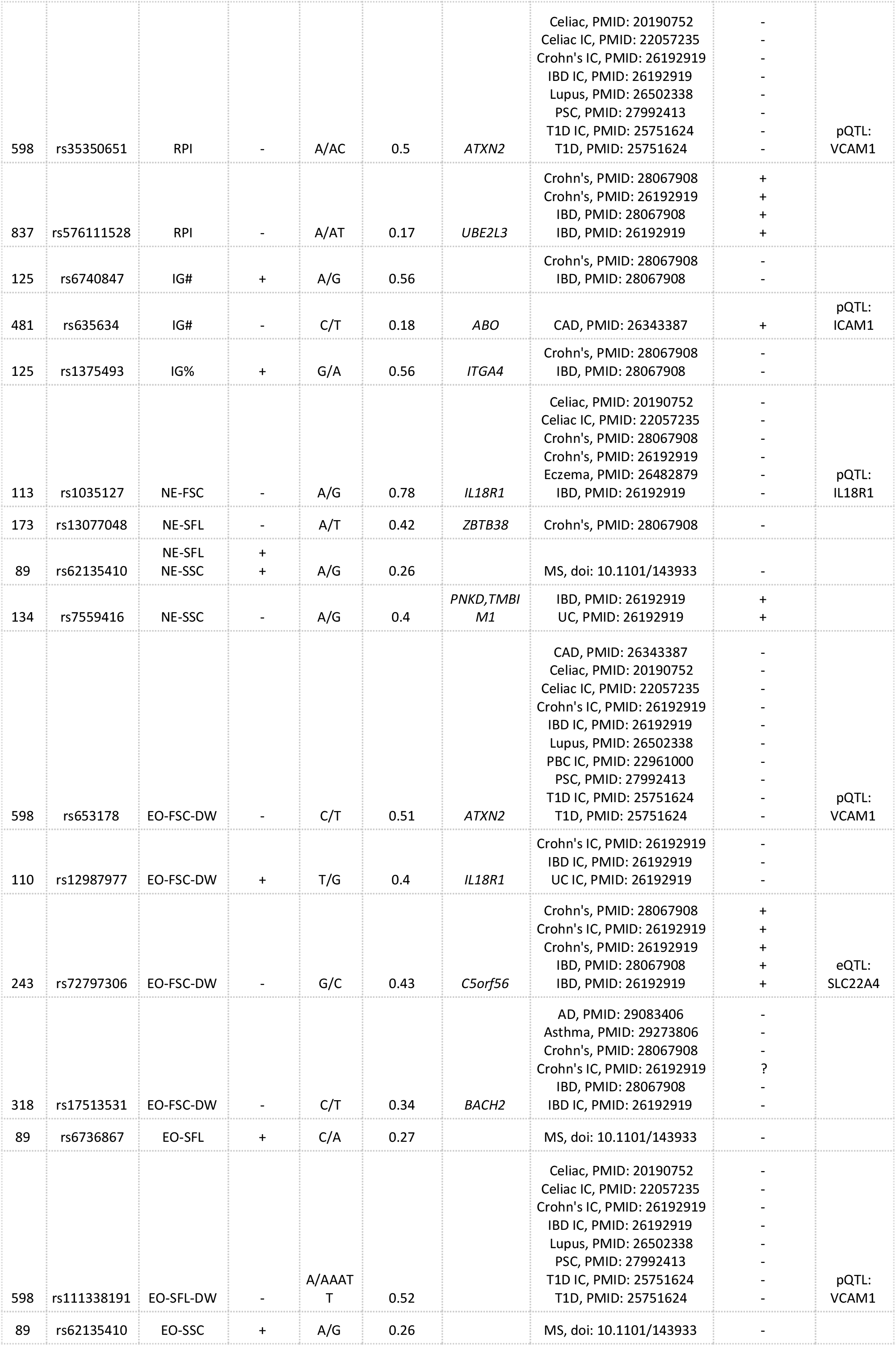
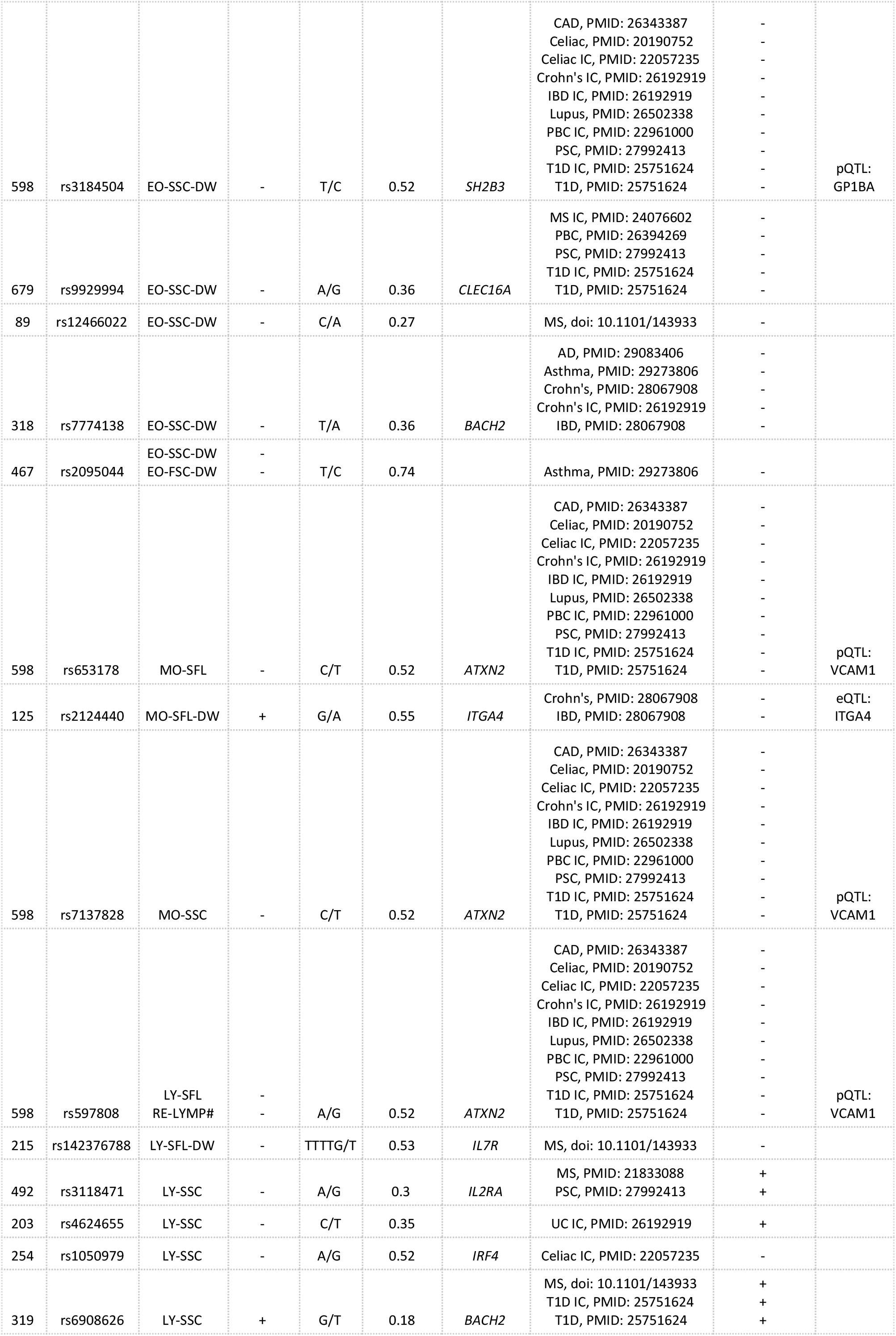
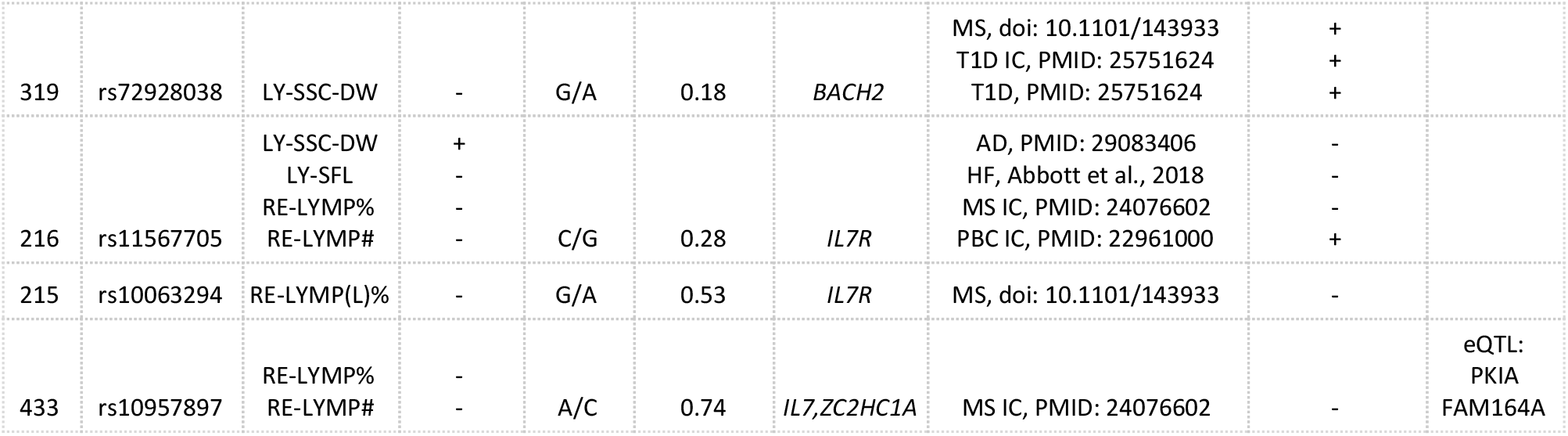
Cytometry Associations that Colocalise with Genetic Associations with Disease Risk. Each row corresponds to a conditionally independent association with cytometry parameters of a cell type and assigned to independent signals by a LD clumping procedure. We also tested for common causal variants between loci of association with Sysmex parameters and disease risk GWAS, blood cell eQTL, and blood plasma pQTL signals. Results show genetic determinants of cytometry parameters have concomitant effects on disease risk.

The ncCBC GWAS identified 153 variant-trait associations (in 101 LD clumps) with lymphocyte traits, of which 15 (in 8 LD clumps) colocalized with at least one genetic association with the risks of multiple sclerosis (MS), celiac disease, primary biliary cirrhosis, hay fever/rhinitis or coronary artery disease (**Table S6**). 12 of these colocalizations (in 5 LD clumps) were with associations for risk of MS, recapitulating the known importance of lymphocytes in the etiology of MS (Legroux and Arbour 2015). The colocalizing associations were located in the transcription factor encoding gene *BACH2*, in the genes encoding receptors for Interleukin(IL)-2 (IL2RA) and IL-7 (IL7R) and in IL-7 itself. IL-2 receptor *α*-chain (the product of IL2RA) is a known therapeutic target for MS. The *IL2RA* colocalising association rs3118471-G (PP=99.9%, AF=30%) corresponds to reduced LY-SSC and an increased risk of MS (International Multiple Sclerosis Genetics Consortium *et al.* 2011). IL2RA is the target of the therapeutic antibody Daclizumab, which is known to be clinically effective in the treatment of MS (Wynn *et al.* 2010; Bielekova *et al.* 2004), but has been withdrawn due to severe side effects including encephalitis (Stork *et al.* 2019; Bielekova *et al.* 2006); (Giovannoni *et al.* 2016; Curto *et al.* 2016; Abbas *et al.* 2011). A second antibody used to treat MS (Natalizumab), which targets the *α*4*β*1 integrin, has been associated with a small number of cases of progressive multifocal leukoencephalopathy (PML), but is still licensed (Bloomgren *et al.* 2012). The risk of developing PML is greater for patients with antibodies against the JC virus.

Alternative therapeutic approaches for MS are required to ameliorate the risk that subsets of patients develop severe side effects on existing drugs. The IL-7 receptor (the product of *IL7R*) is one possible target (Lee *et al.* 2011; Bielekova *et al.* 2006). Preclinical studies in mice show that blockade of IL-7R can ameliorate MS severity (Lawson et al. 2015). Genetic evidence suggests that increasing serum concentration of soluble IL-7R increases the risk of multiple sclerosis (Galarza-Muñoz *et al.* 2017). The colocalising association in *IL7R* with reduced risk of MS is with rs11567705-G (PP=95.7%, AF=27.5%) and the variation in risk is thought to be mediated by a reduction in the soluble isoform of the receptor in favour of the membrane bound isoform (Gregory *et al.* 2007; International Multiple Sclerosis Genetics Consortium *et al.* 2011). Our analysis shows that the association with reduced MS risk colocalizes with an association with decreased RE-LYMP#, a direct measure of the peripheral blood concentration of reactive lymphocytes. Thus our analysis supports the hypothesis that the etiological effect can be explained by the modulation of T-cell activation and provides further support for IL-7R/IL-7 as an efficacious target for MS treatment (Gregory *et al.* 2007).

Three of the 98 variant trait-associations (in 69 LD clumps) with ncCBC phenotypes of monocytes colocalised with disease risk associations, two in *ATXN2* and one in *ITGA4,* all with inflammatory bowel disorder (IBD) associations. An antibody against integrin *α*4**β**7 (encoded by *ITGA4*/*ITGB7*), Vedolizumab, is used to treat IBD (Sandborn *et al.* 2013; Feagan *et al.* 2013). It is assumed to reduce trafficking of *α*4*β*7-positive gut-specific T-helper lymphocyte by diminishing integrin interaction with the mucosal addressin cell adhesion molecule 1 (MadCAM-1) (Soler *et al.* 2009; Rogler 2018). However, a recent study has suggested that the antibody also reduces the ability of monocytes to egress into the colonic mucosa as an alternative mechanism (Rogler 2018; Zeissig *et al.* 2018; Schippers *et al.* 2016). We observed an association in *ITGA4* between rs2124440-A (AF=55.3%) and increased MO-SFL-DW, a measure of the within individual variation of the nucleic acid content of monocytes, and a (sub-critical) association with reduced MO-SFL (*P*-value=2×10^−8^). The MO-SFL association is consistent with the co-localization of the associations with an eQTL decreasing *ITGA4* transcript abundance in monocytes (PP=99.7%). These observations support the hypothesis that the association with IBD in *ITGA4* is partly mediated by monocytes and suggests a non-canonical pathway for the therapeutic effect of Vedolizumab, specifically the inhibition of monocytes.

## DISCUSSION

Over the last decade, GWAS have identified thousands of genetic variants associated with common complex disease risks. However, only a fraction of the biological mechanisms generating these associations are understood, partly because the mediating cell-types and tissues are unknown. For many disease associations blood is a good candidate as a mediating tissue. Blood cells, for instance, are known to play roles in the aetiologies of type 1 diabetes, rheumatoid arthritis, lupus erythematosus, MS, IBD, coronary artery disease and thrombotic stroke. One approach to identifying cell types that might mediate a genetic association with disease is to search for cell type specific associations of the corresponding variant with molecular phenotypes. Colocalization of an eQTL, for example, can suggest a causal gene, cell type and tissue. Unfortunately, molecular intermediate traits such as transcript abundances are expensive to measure and, because it is difficult to sort cells from heterogeneous tissues, their QTLs are often identified from association analysis based on abundance data measured from mixtures of cell types (GTEx Consortium 2015).

We took a complementary approach to molecular phenotyping, using high-throughput flow cytometry to measure 63 ncCBC traits, which capture variation in an array of cell type specific biological processes including transcriptional activity, granule formation, cell degranulation and cell reactivity, in approximately forty thousand healthy blood donors. GWAS analyses of these traits showed that they are heritable, have complex genetic architectures and are affected by variation in genes implicated in a great variety of molecular and cellular pathways perturbing the cellular structures that disturb the laser light as cells pass through a cytometer. One result of this broad biological sensitivity is that cytometry traits have coarser interpretations than intermediate phenotypes of molecular abundance (e.g transcriptomics, proteomics or lipidomics). For example, although cytometry measured ‘neutrophil granularity’ depends on the average number of granules in peripheral blood neutrophils, our results suggest it also depends on the expression of genes corresponding to proteins localized in granules (e.g. *ELANE, MPO, PRTN3*). Consequently, genetic associations with ncCBC cytometry traits can be used in two ways - either as proxies for cis associations with molecular abundance phenotypes, with the target gene/protein infered by physical location, or as a means to identify the key genes regulating the formation and retention of intracellular structures. The latter purpose is exemplified by our multi-omic analysis of the association in *ZFPM2* with PLT-SSC (platelet granularity), which we used to show that the transcription factor FOG2 is a probable regulator of platelet *α*-granularity, mediating variation in the concentrations of a multitude of *α*-granule proteins in the blood plasma. *α*-granule cargo compounds are key to the regulation of thrombus formation, wound healing and thrombus resolution, illustrating that genetic analysis of blood cell cytometry traits can be used to draw inferences about fundamental cellular physiology relevant to cell function.

The integration of our GWAS results with associations from complex disease GWAS demonstrates how ncCBC association signals can capture valuable information about disease etiology, including information useful for drug development. For example, we have shown that genetic associations at particular loci provide evidence supporting the known roles of neutrophils in the etiology of vasculitis, and of lymphocytes in the etiology of MS. We have also shown that the traits can be used to recapitulate the protein and cell-type targets of drugs. Specifically those of Daclizumab (withdrawn due to side effects), which targets the receptor for IL-2 in activated T lymphocytes to treat MS, and those of Vedolizumab which targets the integrin *α*4*β*7, to treat IBD. Finally, we have identified human genetic evidence to support data from previously published studies suggesting that targeting IL-7R to reduce reactive lymphocyte count could be efficacious as a treatment for MS.

## AUTHOR CONTRIBUTIONS

Conceptualization: WHO; Data curation: PA, DV, TJ, KK, RK, LM, JC, LG, JG, SK, SM, JS, KW and WJA; Formal analysis: PA, DV, TJ, EB, LG, DS, JMV and WJA; Funding acquisition: NAW, JD, DJR, EDA, AB, WHO and NS; Investigation: PA, DV, TJ, JS and WJA; Methodology: PA, DV, TJ, SB and WJA; Project administration: NAW, AB, WHO, NS and WJA; Resources: LM, JC, KD, MG, JCK, VS, MF, JP, AB and WHO, NS; Software: PA, DV, TJ, KK, RK, SM, JEP and WJA; Supervision: MF, SB, TK, JEP, AB, WHO, NS and WJA; Visualization: PA, NS and WJA; Writing - original draft: PA, JP, AB, WHO, NS and WJA; Writing-review and editing: PA, DV, EB, LM, JD, DR, VJ, MF, SB, TK, JP, AB, WHO, NS and WJA.

## ACKNOWLEDGMENTS

We are grateful to the participants in the INTERVAL study, whose generosity made this work possible. INTERVAL is a collaboration between the University of Cambridge, the University of Oxford and National Health Service Blood and Transplant (NHSBT; www.nhsbt.nhs.uk). We thank Dr Carmel Moore and the study coordination teams at the University of Cambridge, the University of Oxford and NHSBT. We thank the staff of the Cambridge BioResource and the staff of NHSBT, including the donation staff at the static blood donation centers, for their help with recruitment and for study fieldwork. We thank Mr Mathew Walker and the data management team of the NIHR Blood and Transplant Research Unit in Donor Health and Genomics. We thank Dr Jarob Saker and Dr Joachim Linssen of Sysmex Europe and Mr Stephen Garner, lately retired from the NHSBT Component Development Laboratory, for their invaluable advice on the Sysmex XN-1000 instrument. We thank Joanna Westmoreland of the MRC Laboratory of Molecular Biology for illustration of Figure 1 and 3. We thank members of the Cambridge BioResource Scientific Advisory Board and Management Committee for their support of the study. DNA extraction and genotyping was co-funded by the National Institute for Health Research (NIHR), the NIHR BioResource (http://bioresource.nihr.ac.uk) and the NIHR [Cambridge Biomedical Research Centre at the Cambridge University Hospitals NHS Foundation Trust] [*]. The academic coordinating centre for INTERVAL was supported by core funding from: NIHR Blood and Transplant Research Unit in Donor Health and Genomics (NIHR BTRU-2014-10024), UK Medical Research Council (MR/L003120/1), British Heart Foundation (SP/09/002; RG/13/13/30194; RG/18/13/33946) and the NIHR [Cambridge Biomedical Research Centre at the Cambridge University Hospitals NHS Foundation Trust] [*]. A complete list of the investigators and contributors to the INTERVAL trial is provided in reference [**]. The academic coordinating centre would like to thank blood donor centre staff for their help and to thank blood donors for participating in the INTERVAL trial. This work was supported by Health Data Research UK, which is funded by the UK Medical Research Council (MRC), the Engineering and Physical Sciences Research Council, the Economic and Social Research Council, the Department of Health and Social Care (England), the Chief Scientist Office of the Scottish Government Health and Social Care Directorates, the Health and Social Care Research and Development Division (Welsh Government), the Public Health Agency (Northern Ireland), the British Heart Foundation and the Wellcome Trust. P.A. and D.V. are funded by the National Institute for Health Research Blood and Transplant Research Unit in Donor Health and Genomics (NIHR BTRU-2014-10024). T.J. is funded by a Cambridge University Medical Research Council Doctoral Training Programme and the National Institute for Health Research [Cambridge Biomedical Research Centre at the Cambridge University Hospitals NHS Foundation Trust] [*]. L.M. is supported by a PhD Studentship grant from the Rosetrees Trust. J.C. is supported by a Medical Research Council Clinical Research Training Fellowship. K.D. is supported by Health Education England as an HSST trainee. J.C.K. is funded by a Wellcome Trust Investigator Award (204969/Z/16/Z to J.C.K.), Wellcome Trust Grants (074318 to J.C.K., 090532/Z/09/Z and 203141/Z/16/Z to the Wellcome Centre for Human Genetics core facility), NIHR Oxford Biomedical Research Centre. J.D. is funded by the National Institute for Health Research [Senior Investigator Award] [*]. V.G.S. is supported by NIH grant R01 DK103794, a gift from the Lodish Family to Boston Children’s Hospital, and the New York Stem Cell Foundation (NYSCF). V.G.S. is a NYSCF-Robertson Investigator. M.F. is funded by the British Heart Foundation (FS/18/53/33863). J.E.P. is supported by a UKRI Innovation Fellowship at Health Data Research UK (MR/S004068/1) Research in the W.H.O. laboratory is funded by the International Society on Thrombosis and Haemostasis, the MRC, NHSBT and the Rosetrees Trust. W.J.A. is funded by NHSBT and the National Institute for Health Research [Cambridge Biomedical Research Centre at the Cambridge University Hospitals NHS Foundation Trust] [*]

*The views expressed are those of the authors and not necessarily those of the NHS, the NIHR or the Department of Health and Social Care.

**Di Angelantonio E, Thompson SG, Kaptoge SK, Moore C, Walker M, Armitage J, Ouwehand WH, Roberts DJ, Danesh J, INTERVAL Trial Group. Efficiency and safety of varying the frequency of whole blood donation (INTERVAL): a randomised trial of 45 000 donors. Lancet. 2017 Nov 25;390(10110):2360-2371.

## Supplementary Figure Legends

**Figure S1. Genetic correlation within cytometry parameters.** Heatmap of the genetic correlation calculated between all 63 cytometry parameters, where genetic correlation ranges between −1 and 1 and is calculated from a subset of 63.4 million common variants across the genome in the INTERVAL cohort by LD score regression (Bulik-Sullivan *et al.* 2015). This figure shows limited genetic correlation between parameters of white cells with higher genetic correlation between platelet and red cell parameters and very little correlation between parameters of different cell types. Starred correlations are those insufficiently strong to fall below a Bonferroni significance threshold (*P-*value < 1.28×10^−5^).

**Figure S2. Genetic correlation with clinical full blood count parameters.** Heatmap of the genetic correlation calculated between 63 cytometry parameters and all hematological traits studied by Astle *et al.* (2016) where genetic correlation ranges between −1 and 1 and is calculated from a subset of 63.4 million common variants across the genome in the INTERVAL cohort by LD score regression (Bulik-Sullivan *et al.* 2015). This figure shows limited genetic correlation between cytometry traits and previously studied standard hematological traits. Starred correlations are those insufficiently strong to fall below a Bonferroni significance threshold (*P-*value < 2.27×10^−5^).

**Figure S3. Phenotypic correlation with clinical full blood count parameters.** Heatmap of the Pearson correlation between all 63 cytometry phenotypes and FBC traits (Astle *et al.* 2016) adjusted for technical and environmental covariates, showing limited correlation between cytometry parameters and FBC parameters of white cells and higher correlation for platelet and red cell related parameters.

**Figure S4. Phenotypic correlation within cytometry parameters.** Heatmap of the Pearson correlation between all 63 cytometry phenotypes showing stronger trait correlation adjusted for technical and environmental covariates between phenotypes of red cell and platelet traits than between phenotypes of white cells and low trait correlation between parameters of different cell-types.

**Figure S5. Distribution of LD clumps across cell types.** Distribution of LD clumps across cell types showing limited overlap (31 sets) between LD clumps assigned to different cell types suggesting that the cytometry traits are driven by cell type specific genetic determinants.

**Figure S6. Distribution of LD clumps within a cell type.** Distribution of LD clumps between cytometry traits within a cell type showing limited overlap between the genetic determinants of different cytometry traits (SSC, SFL, FSC, associated distribution width measures, and relevant count measure) of the same cell type. These results show that cytometry traits are influenced by distinct genetic determinants.

**Figure S7. ATAC-seq enrichment analysis for neutrophils, eosinophils, monocytes, and lymphocytes. A-D)** Cell type specific enrichments for FINEMAP 95% credible set variants associated with cytometry parameters. **E)** Legend for panels A-D showing cells from the hematological tree for which specific epigenomic data were assayed. **F)** Diagram of hematological tree corresponding to the hematological cell-type labels displayed in E).

**Figure S8. Distribution of effect sizes for association of rs6993770-T with plasma protein concentration.** We obtained the effect of rs69933770-T on the concentration of 1,472 plasma proteins. The effect of rs6993770-T on plasma concentration was lower on proteins expressed in MKs compared to all proteins and lower still on proteins expressed and localised to *α*-granules. The allelic effect of rs6993770 on plasma proteins localized to *α*-granules is on average 0.369 phenotypic standard deviations lower (*P*-value 8.5×10^−10^), showing that rs6993770-T differentially reduces the plasma concentration of plasma *α*-granule proteins.

## REFERENCES

Abbas, M., P. H. Lalive, M. Chofflon, H-U Simon, C. Chizzolini, and C. Ribi. 2011. “Hypereosinophilia in Patients with Multiple Sclerosis Treated with Natalizumab.” Neurology 77 (16): 1561–64.

Acharya, K. Ravi, and Steven J. Ackerman. 2014. “Eosinophil Granule Proteins: Form and Function.” The Journal of Biological Chemistry 289 (25): 17406–15.

Amulic, Borko, Christel Cazalet, Garret L. Hayes, Kathleen D. Metzler, and Arturo Zychlinsky. 2012. “Neutrophil Function: From Mechanisms to Disease.” Annual Review of Immunology 30 (January): 459–89.

Astle, William J., Heather Elding, Tao Jiang, Dave Allen, Dace Ruklisa, Alice L. Mann, Daniel Mead, et al. 2016. “The Allelic Landscape of Human Blood Cell Trait Variation and Links to Common Complex Disease.” Cell 167 (5): 1415–29.e19.

Attig, Jan, Federico Agostini, Clare Gooding, Anob M. Chakrabarti, Aarti Singh, Nejc Haberman, Julian A. Zagalak, et al. 2018. “Heteromeric RNP Assembly at LINEs Controls Lineage-Specific RNA Processing.” Cell 174 (5): 1067–81.e17.

Becknell, Brian, Tad E. Eichler, Susana Beceiro, Birong Li, Robert S. Easterling, Ashley R. Carpenter, Cindy L. James, et al. 2015. “Ribonucleases 6 and 7 Have Antimicrobial Function in the Human and Murine Urinary Tract.” Kidney International 87 (1): 151–61.

Benner, Christian, Chris C. A. Spencer, Aki S. Havulinna, Veikko Salomaa, Samuli Ripatti, and Matti Pirinen. 2016. “FINEMAP: Efficient Variable Selection Using Summary Data from Genome-Wide Association Studies.” Bioinformatics 32 (10): 1493–1501.

Bielekova, Bibiana, Marta Catalfamo, Susan Reichert-Scrivner, Amy Packer, Magdalena Cerna, Thomas A. Waldmann, Henry McFarland, Pierre A. Henkart, and Roland Martin. 2006. “Regulatory CD56(bright) Natural Killer Cells Mediate Immunomodulatory Effects of IL-2Ralpha-Targeted Therapy (daclizumab) in Multiple Sclerosis.” Proceedings of the National Academy of Sciences of the United States of America 103 (15): 5941–46.

Bielekova, Bibiana, Nancy Richert, Thomas Howard, Gregg Blevins, Silva Markovic-Plese, Jennifer McCartin, Jens Würfel, et al. 2004. “Humanized Anti-CD25 (daclizumab) Inhibits Disease Activity in Multiple Sclerosis Patients Failing to Respond to Interferon *β*.” Proceedings of the National Academy of Sciences of the United States of America 101 (23): 8705–8.

Bloomgren, Gary, Sandra Richman, Christophe Hotermans, Meena Subramanyam, Susan Goelz, Amy Natarajan, Sophia Lee, et al. 2012. “Risk of Natalizumab-Associated Progressive Multifocal Leukoencephalopathy.” The New England Journal of Medicine 366 (20): 1870–80.

Borregaard, Niels, Ole E. Sørensen, and Kim Theilgaard-Mönch. 2007. “Neutrophil Granules: A Library of Innate Immunity Proteins.” Trends in Immunology 28 (8): 340–45.

Bulik-Sullivan, Brendan K., Po-Ru Loh, Hilary K. Finucane, Stephan Ripke, Jian Yang, Schizophrenia Working Group of the Psychiatric Genomics Consortium, Nick Patterson, Mark J. Daly, Alkes L. Price, and Benjamin M. Neale. 2015. “LD Score Regression Distinguishes Confounding from Polygenicity in Genome-Wide Association Studies.” Nature Genetics 47 (3): 291–95.

Calvo-Garrido, Javier, Sergio Carilla-Latorre, Francisco Lázaro-Diéguez, Gustavo Egea, and Ricardo Escalante. 2008. “Vacuole Membrane Protein 1 Is an Endoplasmic Reticulum Protein Required for Organelle Biogenesis, Protein Secretion, and Development.” Molecular Biology of the Cell 19 (8): 3442–53.

Chami, Nathalie, Ming-Huei Chen, Andrew J. Slater, John D. Eicher, Evangelos Evangelou, Salman M. Tajuddin, Latisha Love-Gregory, et al. 2016. “Exome Genotyping Identifies Pleiotropic Variants Associated with Red Blood Cell Traits.” American Journal of Human Genetics 99 (1): 8–21.

Charafeddine, Ali H., Eugenia J. Kim, Dawn M. Maynard, Hong Yi, Timothy A. Weaver, Meral Gunay-Aygun, Maria Russell, William A. Gahl, and Allan D. Kirk. 2012. “Platelet-Derived CD154: Ultrastructural Localization and Clinical Correlation in Organ Transplantation.” American Journal of Transplantation: Official Journal of the American Society of Transplantation and the American Society of Transplant Surgeons 12 (11): 3143–51.

Chatterjee, Madhumita, Zhangsen Huang, Wei Zhang, Lei Jiang, Kjell Hultenby, Linjing Zhu, Hu Hu, Gunnar P. Nilsson, and Nailin Li. 2011. “Distinct Platelet Packaging, Release, and Surface Expression of Proangiogenic and Antiangiogenic Factors on Different Platelet Stimuli.” Blood 117 (14): 3907–11.

Chen, Rui, Ge Jin, and Thomas M. McIntyre. 2017. “The Soluble Protease ADAMDEC1 Released from Activated Platelets Hydrolyzes Platelet Membrane pro-Epidermal Growth Factor (EGF) to Active High-Molecular-Weight EGF.” Journal of Biological Chemistry. https://doi.org/10.1074/jbc.m116.771642.

Cicha, Iwona, Christoph D. Garlichs, Werner G. Daniel, and Margarete Goppelt-Struebe. 2004. “Activated Human Platelets Release Connective Tissue Growth Factor.” Thrombosis and Haemostasis 91 (4): 755–60.

Curto, Elena, Elvira Munteis-Olivas, Eva Balcells, and M. Marisol Domínguez-Álvarez. 2016. “Pulmonary Eosinophilia Associated to Treatment with Natalizumab.” Annals of Thoracic Medicine 11 (3): 224–26.

Deuel, T. F., J. S. Huang, R. T. Proffitt, J. U. Baenziger, D. Chang, and B. B. Kennedy. 1981. “Human Platelet-Derived Growth Factor. Purification and Resolution into Two Active Protein Fractions.” The Journal of Biological Chemistry 256 (17): 8896–99.

Eicher, John D., Nathalie Chami, Tim Kacprowski, Akihiro Nomura, Ming-Huei Chen, Lisa R. Yanek, Salman M. Tajuddin, et al. 2016. “Platelet-Related Variants Identified by Exomechip Meta-Analysis in 157,293 Individuals.” American Journal of Human Genetics 99 (1): 40–55.

Fang, Li, Yibing Yan, Laszlo G. Komuves, Shirlee Yonkovich, Carol M. Sullivan, Bradley Stringer, Sarah Galbraith, et al. 2004. “PDGF C Is a Selective Alpha Platelet-Derived Growth Factor Receptor Agonist That Is Highly Expressed in Platelet Alpha Granules and Vascular Smooth Muscle.” Arteriosclerosis, Thrombosis, and Vascular Biology 24 (4): 787–92.

Feagan, Brian G., Paul Rutgeerts, Bruce E. Sands, Stephen Hanauer, Jean-Frédéric Colombel, William J. Sandborn, Gert Van Assche, et al. 2013. “Vedolizumab as Induction and Maintenance Therapy for Ulcerative Colitis.” The New England Journal of Medicine 369 (8): 699–710.

Galarza-Muñoz, Gaddiel, Farren B. S. Briggs, Irina Evsyukova, Geraldine Schott-Lerner, Edward M. Kennedy, Tinashe Nyanhete, Liuyang Wang, et al. 2017. “Human Epistatic Interaction Controls IL7R Splicing and Increases Multiple Sclerosis Risk.” Cell 169 (1): 72–84.e13.

Gieger, Christian, Aparna Radhakrishnan, Ana Cvejic, Weihong Tang, Eleonora Porcu, Giorgio Pistis, Jovana Serbanovic-Canic, et al. 2011. “New Gene Functions in Megakaryopoiesis and Platelet Formation.” Nature 480 (7376): 201–8.

Giovannoni, Gavin, Ludwig Kappos, Ralf Gold, Bhupendra O. Khatri, Krzysztof Selmaj, Kimberly Umans, Steven J. Greenberg, Marianne Sweetser, Jacob Elkins, and Peter McCroskery. 2016. “Safety and Tolerability Profile of Daclizumab in Patients with Relapsing-Remitting Multiple Sclerosis: An Integrated Analysis of Clinical Studies.” Multiple Sclerosis and Related Disorders 9 (September): 36–46.

Grassi, Luigi, Osagie G. Izuogu, Natasha A. N. Jorge, Denis Seyres, Mariona Bustamante, Frances Burden, Samantha Farrow, et al. 2019. “Cell Type Specific Novel lincRNAs and circRNAs in the BLUEPRINT Haematopoietic Transcriptomes Atlas.” bioRxiv. https://doi.org/10.1101/764613.

Grassi, Luigi, Farzin Pourfarzad, Sebastian Ullrich, Angelika Merkel, Felipe Were, Enrique Carrillo-de-Santa-Pau, Guoqiang Yi, et al. 2018. “Dynamics of Transcription Regulation in Human Bone Marrow Myeloid Differentiation to Mature Blood Neutrophils.” Cell Reports 24 (10): 2784–94.

Gregory, Simon G., Silke Schmidt, Puneet Seth, Jorge R. Oksenberg, John Hart, Angela Prokop, Stacy J. Caillier, et al. 2007. “Interleukin 7 Receptor Alpha Chain (IL7R) Shows Allelic and Functional Association with Multiple Sclerosis.” Nature Genetics 39 (9): 1083–91.

GTEx Consortium. 2015. “Human Genomics. The Genotype-Tissue Expression (GTEx) Pilot Analysis: Multitissue Gene Regulation in Humans.” Science 348 (6235): 648–60.

Guo, Jia, Mengjiao Zhou, Xin Liu, Yunzhi Pan, Runjun Yang, Zhihui Zhao, and Boxing Sun. 2018. “Porcine IFI30 Inhibits PRRSV Proliferation and Host Cell Apoptosis in Vitro.” Gene 649 (April): 93–98.

Han, G. Celine, Vinesh Vinayachandran, Alain R. Bataille, Bongsoo Park, Ka Yim Chan-Salis, Cheryl A. Keller, Maria Long, Shaun Mahony, Ross C. Hardison, and B. Franklin Pugh. 2016. “Genome-Wide Organization of GATA1 and TAL1 Determined at High Resolution.” Molecular and Cellular Biology 36 (1): 157–72.

Henriot, I., E. Launay, M. Boubaya, L. Cremet, M. Illiaquer, H. Caillon, A. Desjonquères, B. Gillet, M. C. Béné, and M. Eveillard. 2017. “New Parameters on the Hematology Analyzer XN-10 (SysmexTM) Allow to Distinguish Childhood Bacterial and Viral Infections.” International Journal of Laboratory Hematology 39 (1): 14–20.

Holten, Thijs C. van, Onno B. Bleijerveld, Patrick Wijten, Philip G. de Groot, Albert J. R. Heck, Arjan D. Barendrecht, Tesy H. Merkx, Arjen Scholten, and Mark Roest. 2014. “Quantitative Proteomics Analysis Reveals Similar Release Profiles Following Specific PAR-1 or PAR-4 Stimulation of Platelets.” Cardiovascular Research 103 (1): 140–46.

Huang, C-L, C.-L. Huang, J.-C. Cheng, A. Stern, J.-T. Hsieh, C.-H. Liao, and C.-P. Tseng. 2006. “Disabled-2 Is a Novel IIb-Integrin-Binding Protein That Negatively Regulates Platelet-Fibrinogen Interactions and Platelet Aggregation.” Journal of Cell Science. https://doi.org/10.1242/jcs.03195.

International Multiple Sclerosis Genetics Consortium, Wellcome Trust Case Control Consortium 2, Stephen Sawcer, Garrett Hellenthal, Matti Pirinen, Chris C. A. Spencer, Nikolaos A. Patsopoulos, et al. 2011. “Genetic Risk and a Primary Role for Cell-Mediated Immune Mechanisms in Multiple Sclerosis.” Nature 476 (7359): 214–19.

Johnson, Andrew D., Lisa R. Yanek, Ming-Huei Chen, Nauder Faraday, Martin G. Larson, Geoffrey Tofler, Shiow J. Lin, et al. 2010. “Genome-Wide Meta-Analyses Identifies Seven Loci Associated with Platelet Aggregation in Response to Agonists.” Nature Genetics 42 (7): 608–13.

Kaplan, D. R., F. C. Chao, C. D. Stiles, H. N. Antoniades, and C. D. Scher. 1979. “Platelet Alpha Granules Contain a Growth Factor for Fibroblasts.” Blood 53 (6): 1043–52.

Kassouf, Mira T., Jim R. Hughes, Stephen Taylor, Simon J. McGowan, Shamit Soneji, Angela L. Green, Paresh Vyas, and Catherine Porcher. 2010. “Genome-Wide Identification of TAL1’s Functional Targets: Insights into Its Mechanisms of Action in Primary Erythroid Cells.” Genome Research 20 (8): 1064–83.

Klarin, Derek, Emma Busenkell, Renae Judy, Julie Lynch, Michael Levin, Jeffery Haessler, Krishna Aragam, et al. 2019. “Genome-Wide Association Analysis of Venous Thromboembolism Identifies New Risk Loci and Genetic Overlap with Arterial Vascular Disease.” Nature Genetics 51 (11): 1574–79.

Klarin, Derek, Connor A. Emdin, Pradeep Natarajan, Mark F. Conrad, INVENT Consortium, and Sekar Kathiresan. 2017. “Genetic Analysis of Venous Thromboembolism in UK Biobank Identifies the ZFPM2 Locus and Implicates Obesity as a Causal Risk Factor.” Circulation. Cardiovascular Genetics 10 (2). https://doi.org/10.1161/CIRCGENETICS.116.001643.

Kreuzhuber, Roman. 2018. “The Effect of Non-Coding Variants on Gene Transcription in Human Blood Cell Types,” November.

Kundu, Kousik, Alice L. Mann, Manuel Tardaguila, Stephen Watt, Hannes Ponstingl, Louella Vasquez, Nicholas W. Morrell, et al. 2020. “Genetic Associations at Regulatory Phenotypes Improve Fine-Mapping of Causal Variants for Twelve Immune-Mediated Diseases.” bioRxiv. https://doi.org/10.1101/2020.01.15.907436.

Lawson, Brian R., Rosana Gonzalez-Quintial, Theodoros Eleftheriadis, Michael A. Farrar, Stephen D. Miller, Karsten Sauer, Dorian B. McGavern, Dwight H. Kono, Roberto Baccala, and Argyrios N. Theofilopoulos. 2015. “Interleukin-7 Is Required for CD4(+) T Cell Activation and Autoimmune Neuroinflammation.” Clinical Immunology 161 (2): 260–69.

Lee, Li-Fen, Robert Axtell, Guang Huan Tu, Kathryn Logronio, Jeanette Dilley, Jessica Yu, Mathias Rickert, et al. 2011. “IL-7 Promotes T(H)1 Development and Serum IL-7 Predicts Clinical Response to Interferon-*β* in Multiple Sclerosis.” Science Translational Medicine 3 (93): 93ra68.

Legroux, Laurine, and Nathalie Arbour. 2015. “Multiple Sclerosis and T Lymphocytes: An Entangled Story.” Journal of Neuroimmune Pharmacology: The Official Journal of the Society on NeuroImmune Pharmacology 10 (4): 528–46.

Linssen, J., S. Aderhold, A. Nierhaus, D. Frings, C. Kaltschmidt, and K. Zänker. 2008. “Automation and Validation of a Rapid Method to Assess Neutrophil and Monocyte Activation by Routine Fluorescence Flow Cytometry in Vitro.” Cytometry. Part B, Clinical Cytometry 74 (5): 295–309.

Loh, Po-Ru, George Tucker, Brendan K. Bulik-Sullivan, Bjarni J. Vilhjálmsson, Hilary K. Finucane, Rany M. Salem, Daniel I. Chasman, et al. 2015. “Efficient Bayesian Mixed-Model Analysis Increases Association Power in Large Cohorts.” Nature Genetics 47 (3): 284–90.

MacGlashan, Donald W., Jr. 2013. “Basophil Activation Testing.” The Journal of Allergy and Clinical Immunology 132 (4): 777–87.

Maynard, Dawn M., Harry F. G. Heijnen, William A. Gahl, and Meral Gunay-Aygun. 2010. “The *α*-Granule Proteome: Novel Proteins in Normal and Ghost Granules in Gray Platelet Syndrome.” Journal of Thrombosis and Haemostasis: JTH 8 (8): 1786–96.

Maynard, D. M., H. F. G. Heijnen, M. K. Horne, J. G. White, and W. A. Gahl. 2007. “Proteomic Analysis of Platelet Alpha-Granules Using Mass Spectrometry.” Journal of Thrombosis and Haemostasis: JTH 5 (9): 1945–55.

McLaren, William, Laurent Gil, Sarah E. Hunt, Harpreet Singh Riat, Graham R. S. Ritchie, Anja Thormann, Paul Flicek, and Fiona Cunningham. 2016. “The Ensembl Variant Effect Predictor.” Genome Biology 17 (1): 122.

Merkel, Peter A., Gang Xie, Paul A. Monach, Xuemei Ji, Dominic J. Ciavatta, Jinyoung Byun, Benjamin D. Pinder, et al. 2017. “Identification of Functional and Expression Polymorphisms Associated With Risk for Antineutrophil Cytoplasmic Autoantibody–Associated Vasculitis.” Arthritis & Rheumatology 69 (5): 1054–66.

Moore, Carmel, Jennifer Sambrook, Matthew Walker, Zoe Tolkien, Stephen Kaptoge, David Allen, Susan Mehenny, et al. 2014. “The INTERVAL Trial to Determine Whether Intervals between Blood Donations Can Be Safely and Acceptably Decreased to Optimise Blood Supply: Study Protocol for a Randomised Controlled Trial.” Trials 15 (September): 363.

Moreau, Thomas, Amanda L. Evans, Louella Vasquez, Marloes R. Tijssen, Ying Yan, Matthew W. Trotter, Daniel Howard, et al. 2016. “Large-Scale Production of Megakaryocytes from Human Pluripotent Stem Cells by Chemically Defined Forward Programming.” Nature Communications 7 (April): 11208.

Parsons, Martin E. M., Paulina B. Szklanna, Jose A. Guerrero, Kieran Wynne, Feidhlim Dervin, Karen O’Connell, Seamus Allen, et al. 2018. “Platelet Releasate Proteome Profiling Reveals a Core Set of Proteins with Low Variance between Healthy Adults.” Proteomics 18 (15): e1800219.

Petersen, Romina, John J. Lambourne, Biola M. Javierre, Luigi Grassi, Roman Kreuzhuber, Dace Ruklisa, Isabel M. Rosa, et al. 2017. “Platelet Function Is Modified by Common Sequence Variation in Megakaryocyte Super Enhancers.” Nature Communications 8 (July): 16058.

Pickrell, Joseph K., Tomaz Berisa, Jimmy Z. Liu, Laure Ségurel, Joyce Y. Tung, and David A. Hinds. 2016. “Detection and Interpretation of Shared Genetic Influences on 42 Human Traits.” Nature Genetics 48 (7): 709–17.

Pruim, Randall J., Ryan P. Welch, Serena Sanna, Tanya M. Teslovich, Peter S. Chines, Terry P. Gliedt, Michael Boehnke, Gonçalo R. Abecasis, and Cristen J. Willer. 2010. “LocusZoom: Regional Visualization of Genome-Wide Association Scan Results.” Bioinformatics 26 (18): 2336–37.

Rogler, Gerhard. 2018. “Mechanism of Action of Vedolizumab: Do We Really Understand It?” Gut.

Sandborn, William J., Brian G. Feagan, Paul Rutgeerts, Stephen Hanauer, Jean-Frédéric Colombel, Bruce E. Sands, Milan Lukas, et al. 2013. “Vedolizumab as Induction and Maintenance Therapy for Crohn’s Disease.” The New England Journal of Medicine 369 (8): 711–21.

Schippers, A., M. Muschaweck, T. Clahsen, S. Tautorat, L. Grieb, K. Tenbrock, N. Gaßler, and N. Wagner. 2016. “*β*7-Integrin Exacerbates Experimental DSS-Induced Colitis in Mice by Directing Inflammatory Monocytes into the Colon.” Mucosal Immunology 9 (2): 527–38.

Soler, Dulce, Tobias Chapman, Li-Li Yang, Tim Wyant, Robert Egan, and Eric R. Fedyk. 2009. “The Binding Specificity and Selective Antagonism of Vedolizumab, an Anti-alpha4beta7 Integrin Therapeutic Antibody in Development for Inflammatory Bowel Diseases.” The Journal of Pharmacology and Experimental Therapeutics 330 (3): 864–75.

Stork, Lidia, Wolfgang Brück, Phillip von Gottberg, Ulrich Pulkowski, Florian Kirsten, Markus Glatzel, Sebastian Rauer, et al. 2019. “Severe Meningo-/encephalitis after Daclizumab Therapy for Multiple Sclerosis.” Multiple Sclerosis 25 (12): 1618–32.

Sun, Benjamin B., Joseph C. Maranville, James E. Peters, David Stacey, James R. Staley, James Blackshaw, Stephen Burgess, et al. 2018. “Genomic Atlas of the Human Plasma Proteome.” Nature 558 (7708): 73–79.

Sysmex. 2014. “Automated Hematology Analyzer XN Series (XN-1000) Instructions for Use.” Sysmex Corporation, Kobe, Japan, February.

Tajuddin, Salman M., Ursula M. Schick, John D. Eicher, Nathalie Chami, Ayush Giri, Jennifer A. Brody, W. David Hill, et al. 2016. “Large-Scale Exome-Wide Association Analysis Identifies Loci for White Blood Cell Traits and Pleiotropy with Immune-Mediated Diseases.” American Journal of Human Genetics 99 (1): 22–39.

Tamura, Shogo, Hidenori Suzuki, Yuji Hirowatari, Masanao Hatase, Ayumi Nagasawa, Kazuhiko Matsuno, Seiichi Kobayashi, and Takanori Moriyama. 2011. “Release Reaction of Brain-Derived Neurotrophic Factor (BDNF) through PAR1 Activation and Its Two Distinct Pools in Human Platelets.” Thrombosis Research 128 (5): e55–61.

The Gene Ontology Consortium. 2019. “The Gene Ontology Resource: 20 Years and Still GOing Strong.” Nucleic Acids Research 47 (D1): D330–38.

Ulirsch, Jacob C., Caleb A. Lareau, Erik L. Bao, Leif S. Ludwig, Michael H. Guo, Christian Benner, Ansuman T. Satpathy, et al. 2019. “Interrogation of Human Hematopoiesis at Single-Cell and Single-Variant Resolution.” Nature Genetics 51 (4): 683–93.

Wang, Yuzhi, Xue Lin, Xue Gong, Lele Wu, Jun Zhang, Weiguang Liu, Jian Li, and Liming Chen. 2018. “Atypical GATA Transcription Factor TRPS1 Represses Gene Expression by Recruiting CHD4/NuRD(MTA2) and Suppresses Cell Migration and Invasion by Repressing TP63 Expression.” Oncogenesis 7 (12): 96.

West, Laura Ciaccia, and Peter Cresswell. 2013. “Expanding Roles for GILT in Immunity.” Current Opinion in Immunology 25 (1): 103–8.

Wijten, Patrick, Thijs van Holten, Liy Liy Woo, Onno B. Bleijerveld, Mark Roest, Albert J. R. Heck, and Arjen Scholten. 2013. “High Precision Platelet Releasate Definition by Quantitative Reversed Protein Profiling--Brief Report.” Arteriosclerosis, Thrombosis, and Vascular Biology 33 (7): 1635–38.

Wynn, Daniel, Michael Kaufman, Xavier Montalban, Timothy Vollmer, Jack Simon, Jacob Elkins, Gilmore O’Neill, et al. 2010. “Daclizumab in Active Relapsing Multiple Sclerosis (CHOICE Study): A Phase 2, Randomised, Double-Blind, Placebo-Controlled, Add-on Trial with Interferon Beta.” Lancet Neurology 9 (4): 381–90.

Xu, Changjiang, Ioanna Tachmazidou, Klaudia Walter, Antonio Ciampi, Eleftheria Zeggini, and Celia M. T. Greenwood. 2014. “Estimating Genome-Wide Significance for Whole-Genome Sequencing Studies.” Genetic Epidemiology 38 (4): 281–90.

Zeissig, Sebastian, Elisa Rosati, C. Marie Dowds, Konrad Aden, Johannes Bethge, Berenice Schulte, Wei Hung Pan, et al. 2018. “Vedolizumab Is Associated with Changes in Innate rather than Adaptive Immunity in Patients with Inflammatory Bowel Disease.” Gut 68 (1): 25–39.

Zhang, Yu, Lin An, Feng Yue, and Ross C. Hardison. 2016. “Jointly Characterizing Epigenetic Dynamics across Multiple Human Cell Types.” Nucleic Acids Research 44 (14): 6721–31.

Zhao, Yan G., Yong Chen, Guangyan Miao, Hongyu Zhao, Wenyan Qu, Dongfang Li, Zheng Wang, et al. 2017. “The ER-Localized Transmembrane Protein EPG-3/VMP1 Regulates SERCA Activity to Control ER-Isolation Membrane Contacts for Autophagosome Formation.” Molecular Cell 67 (6): 974–89.e6.

Zimmermann, Mathias, Malte Cremer, Christina Hoffmann, Karin Weimann, and Andreas Weimann. 2011. “Granularity Index of the SYSMEX XE-5000 Hematology Analyzer as a Replacement for Manual Microscopy of Toxic Granulation Neutrophils in Patients with Inflammatory Diseases.” Clinical Chemistry and Laboratory Medicine: CCLM / FESCC 49 (7): 1193–98.

Zufferey, Anne, Domitille Schvartz, Séverine Nolli, Jean-Luc Reny, Jean-Charles Sanchez, and Pierre Fontana. 2014. “Characterization of the Platelet Granule Proteome: Evidence of the Presence of MHC1 in Alpha-Granules.” Journal of Proteomics 101 (April): 130–40.

