## Supplementary Figures for "Genetic Analyses of Blood Cell Structure for Biological and Pharmacological Inference"

supplementary_figure_1 2
supplementary_figure_2 3
supplementary_figure_3 4
supplementary_figure_4 5
supplementary_figure_5 6
supplementary_figure_6 7
supplementary_figure_7 8
supplementary_figure_8 9

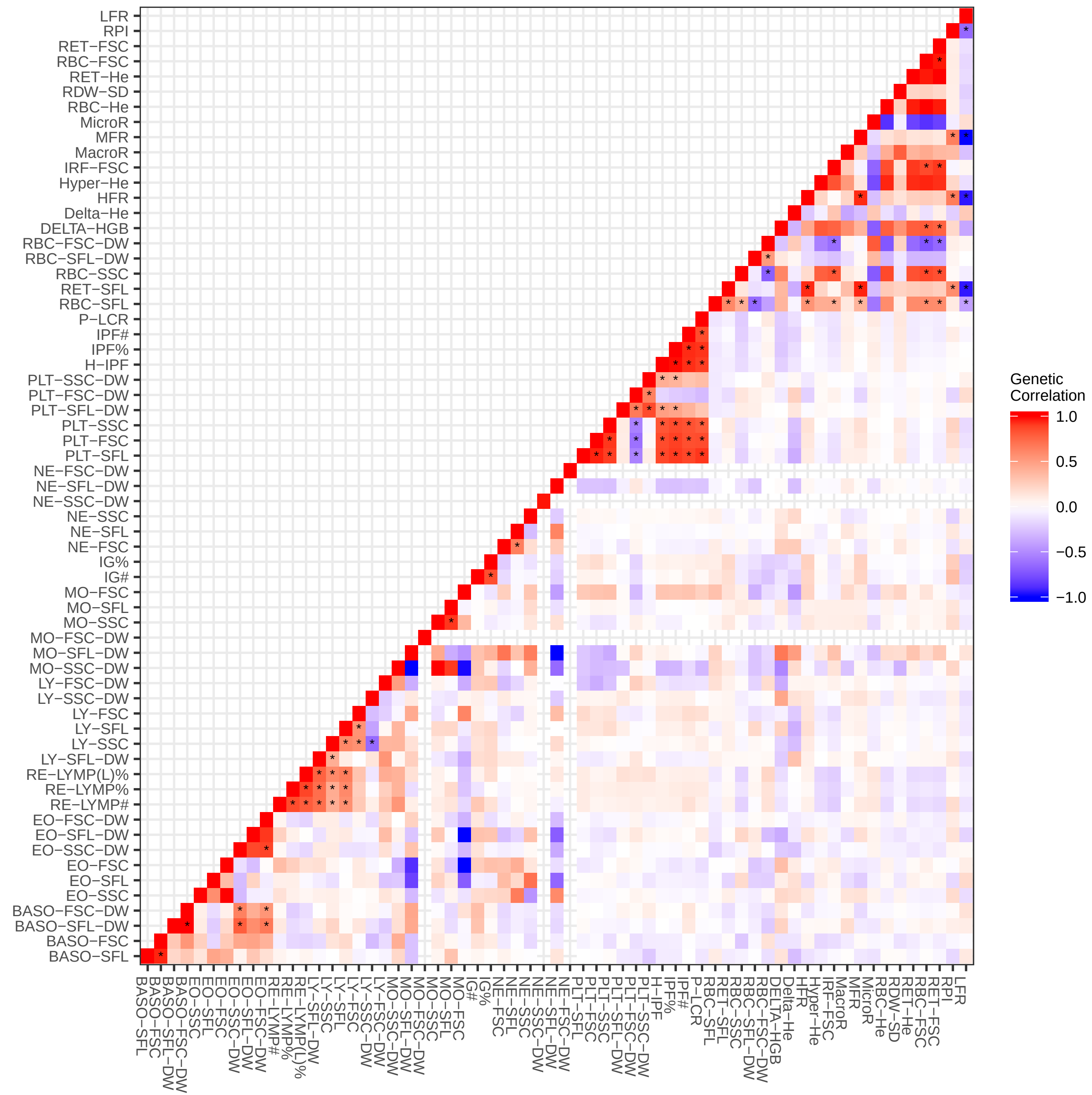
LFR
RPI
RET−FSC
RBC−FSC
RET−He
RDW−SD
RBC−He
MicroR
MFR
MacroR
IRF−FSC
Hyper−He
HFR
Delta−He
DELTA−HGB
RBC−FSC−DW
RBC−SFL−DW
RBC−SSC
RET−SFL
RBC−SFL
P−LCR
IPF#
IPF%
H−IPF
PLT−SSC−DW
PLT−FSC−DW
PLT−SFL−DW
PLT−SSC
PLT−FSC
PLT−SFL
NE−FSC−DW
NE−SFL−DW
NE−SSC−DW
NE−SSC
NE−SFL
NE−FSC
IG%
IG#
MO−FSC
MO−SFL
MO−SSC
MO−FSC−DW
MO−SFL−DW
MO−SSC−DW
LY−FSC−DW
LY−SSC−DW
LY−FSC
LY−SFL
LY−SSC
LY−SFL−DW
RE−LYMP(L)%
RE−LYMP%
RE−LYMP#
EO−FSC−DW
EO−SFL−DW
EO−SSC−DW
EO−FSC
EO−SFL
EO−SSC
BASO−FSC−DW
BASO−SFL−DW
BASO−FSC
BASO−SFL
Genetic Correlation
1.0
0.5
0.0
−0.5
−1.0

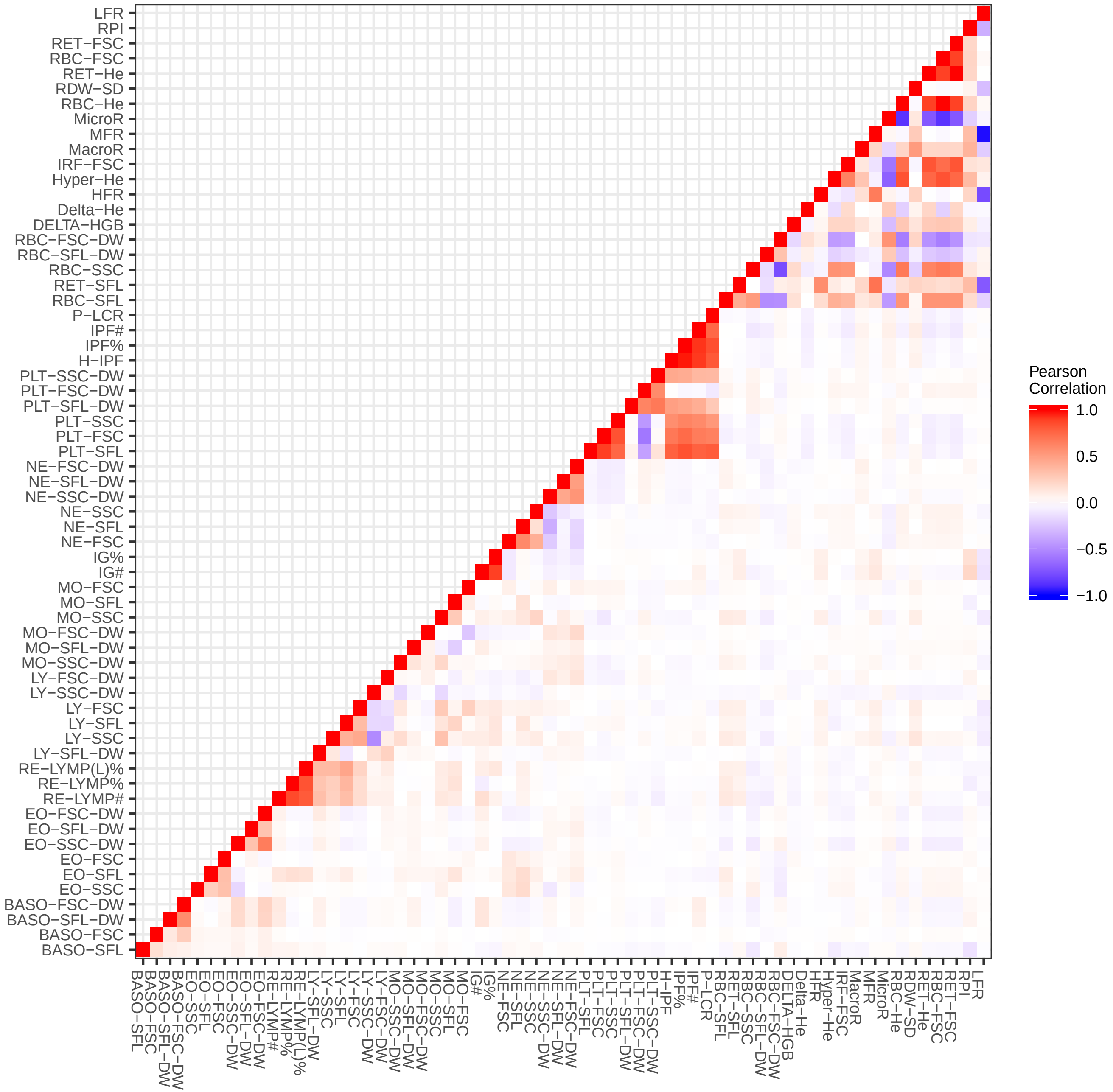
LFR
RPI
RET−FSC
RBC−FSC
RET−He
RDW−SD
RBC−He
MicroR
MFR
MacroR
IRF−FSC
Hyper−He
HFR
Delta−He
DELTA−HGB
RBC−FSC−DW
RBC−SFL−DW
RBC−SSC
RET−SFL
RBC−SFL
P−LCR
IPF#
IPF%
H−IPF
PLT−SSC−DW
PLT−FSC−DW
PLT−SFL−DW
PLT−SSC
PLT−FSC
PLT−SFL
NE−FSC−DW
NE−SFL−DW
NE−SSC−DW
NE−SSC
NE−SFL
NE−FSC
IG%
IG#
MO−FSC
MO−SFL
MO−SSC
MO−FSC−DW
MO−SFL−DW
MO−SSC−DW
LY−FSC−DW
LY−SSC−DW
LY−FSC
LY−SFL
LY−SSC
LY−SFL−DW
RE−LYMP(L)%
RE−LYMP%
RE−LYMP#
EO−FSC−DW
EO−SFL−DW
EO−SSC−DW
EO−FSC
EO−SFL
EO−SSC
BASO−FSC−DW
BASO−SFL−DW
BASO−FSC
BASO−SFL
Pearson Correlation
1.0
0.5
0.0
−0.5
−1.0

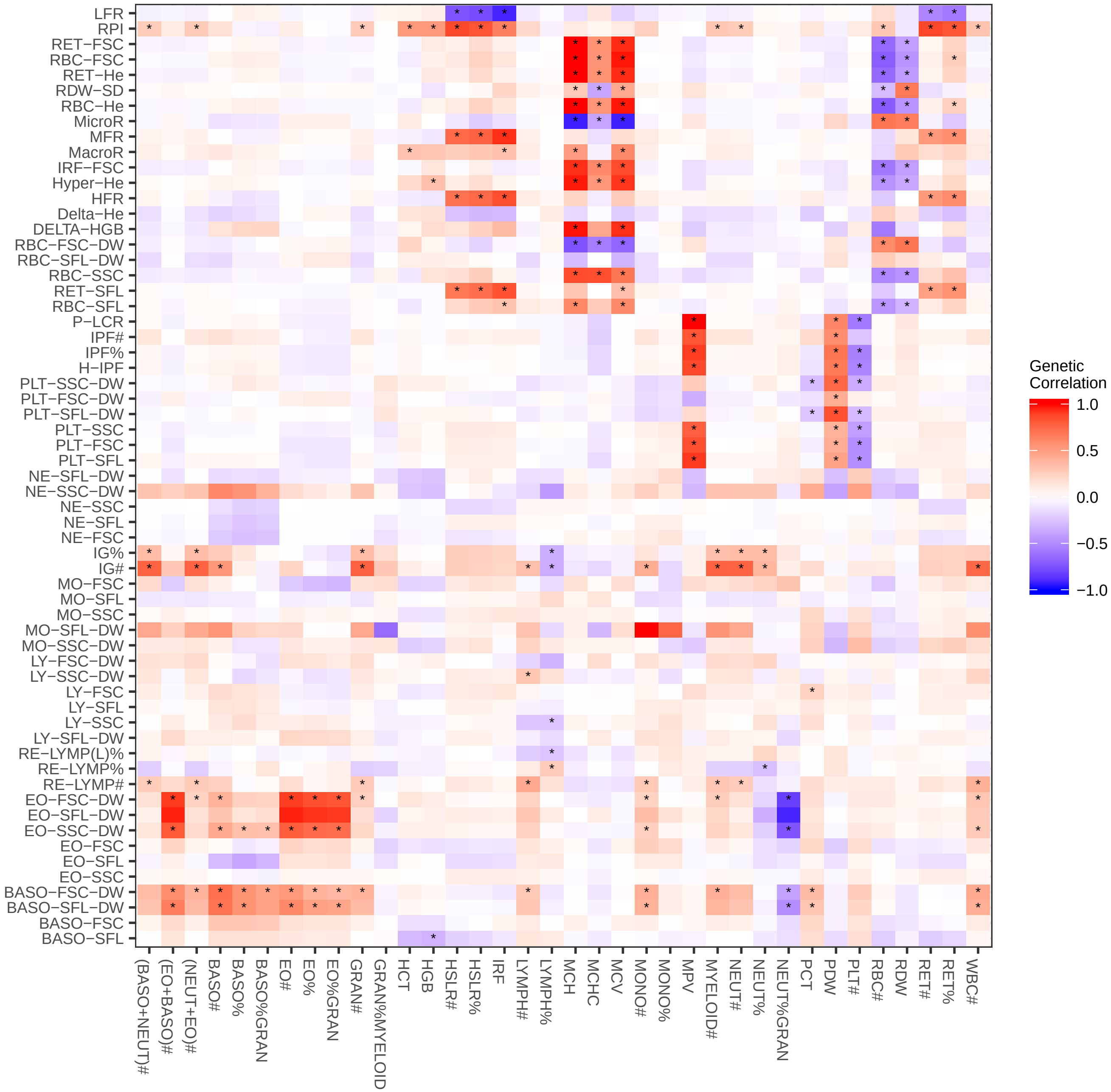

LFR
RPI
RET−FSC
RBC−FSC
RET−He
RDW−SD
RBC−He
MicroR
MFR
MacroR
IRF−FSC
Hyper−He
HFR
Delta−He
DELTA−HGB
RBC−FSC−DW
RBC−SFL−DW
RBC−SSC
RET−SFL
RBC−SFL
P−LCR
IPF#
IPF%
H−IPF
PLT−SSC−DW
PLT−FSC−DW
PLT−SFL−DW
PLT−SSC
PLT−FSC
PLT−SFL
NE−SFL−DW
NE−SSC−DW
NE−SSC
NE−SFL
NE−FSC
IG%
IG#
MO−FSC
MO−SFL
MO−SSC
MO−SFL−DW
MO−SSC−DW
LY−FSC−DW
LY−SSC−DW
LY−FSC
LY−SFL
LY−SSC
LY−SFL−DW
RE−LYMP(L)%
RE−LYMP%
RE−LYMP#
EO−FSC−DW
EO−SFL−DW
EO−SSC−DW
EO−FSC
EO−SFL
EO−SSC
BASO−FSC−DW
BASO−SFL−DW
BASO−FSC
BASO−SFL
(BASO+NEUT)#
(EO+BASO)#
(NEUT+EO)#
BASO#
BASO%
BASO%GRAN
EO#
EO%
EO%GRAN
GRAN#
GRAN%MYELOID
HCT
HGB
HSLR#
HSLR%
IRF
LYMPH#
LYMPH%
MCH
MCHC
MCV
MONO#
MONO%
MPV
MYELOID#
NEUT#
NEUT%
NEUT%GRAN
PCT
PDW
PLT#
RBC#
RDW
RET#
RET%
WBC#
Genetic Correlation
1.0
0.5
0.0
−0.5
−1.0

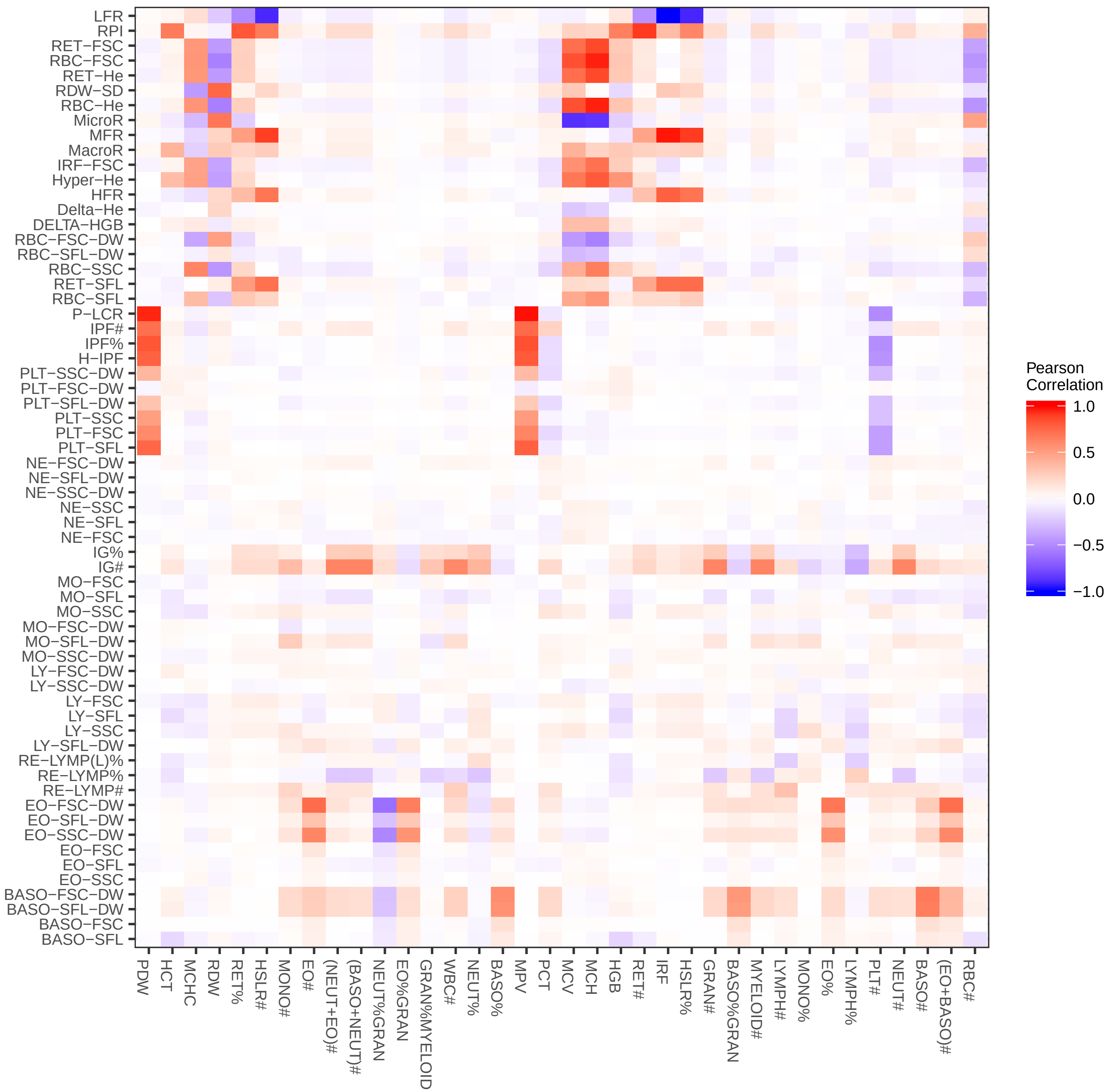

LFR
RPI
RET−FSC
RBC−FSC
RET−He
RDW−SD
RBC−He
MicroR
MFR
MacroR
IRF−FSC
Hyper−He
HFR
Delta−He
DELTA−HGB
RBC−FSC−DW
RBC−SFL−DW
RBC−SSC
RET−SFL
RBC−SFL
P−LCR
IPF#
IPF%
H−IPF
PLT−SSC−DW
PLT−FSC−DW
PLT−SFL−DW
PLT−SSC
PLT−FSC
PLT−SFL
NE−FSC−DW
NE−SFL−DW
NE−SSC−DW
NE−SSC
NE−SFL
NE−FSC
IG%
IG#
MO−FSC
MO−SFL
MO−SSC
MO−FSC−DW
MO−SFL−DW
MO−SSC−DW
LY−FSC−DW
LY−SSC−DW
LY−FSC
LY−SFL
LY−SSC
LY−SFL−DW
RE−LYMP(L)%
RE−LYMP%
RE−LYMP#
EO−FSC−DW
EO−SFL−DW
EO−SSC−DW
EO−FSC
EO−SFL
EO−SSC
BASO−FSC−DW
BASO−SFL−DW
BASO−FSC
BASO−SFL
PDW
HCT
MCHC
RDW
RET%
HSLR#
MONO#
EO#
(NEUT+EO)#
(BASO+NEUT)#
NEUT%GRAN
EO%GRAN
GRAN%MYELOID
WBC#
NEUT%
BASO%
MPV
PCT
MCV
MCH
HGB
RET#
IRF
HSLR%
GRAN#
BASO%GRAN
MYELOID#
LYMPH#
MONO%
EO%
LYMPH%
PLT#
NEUT#
BASO#
(EO+BASO)#
RBC#
Pearson Correlation
1.0
0.5
0.0
−0.5
−1.0

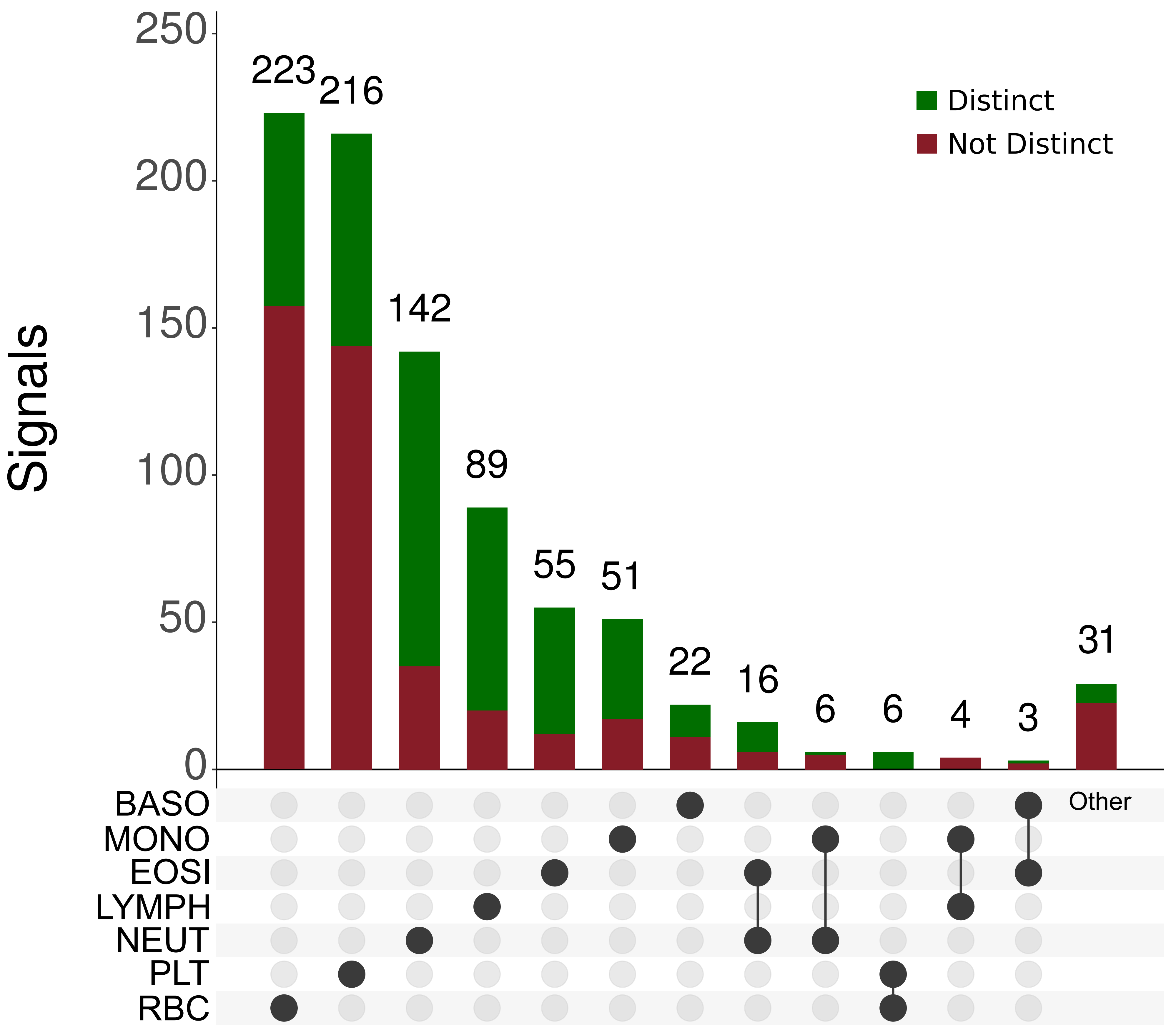
Signals
250
200
150
100
50
0
223
216
142
89
55
51
22
16
6
6
4
3
31
Distinct
Not Distinct
BASO
MONO
EOSI
LYMPH
NEUT
PLT
RBC
Other

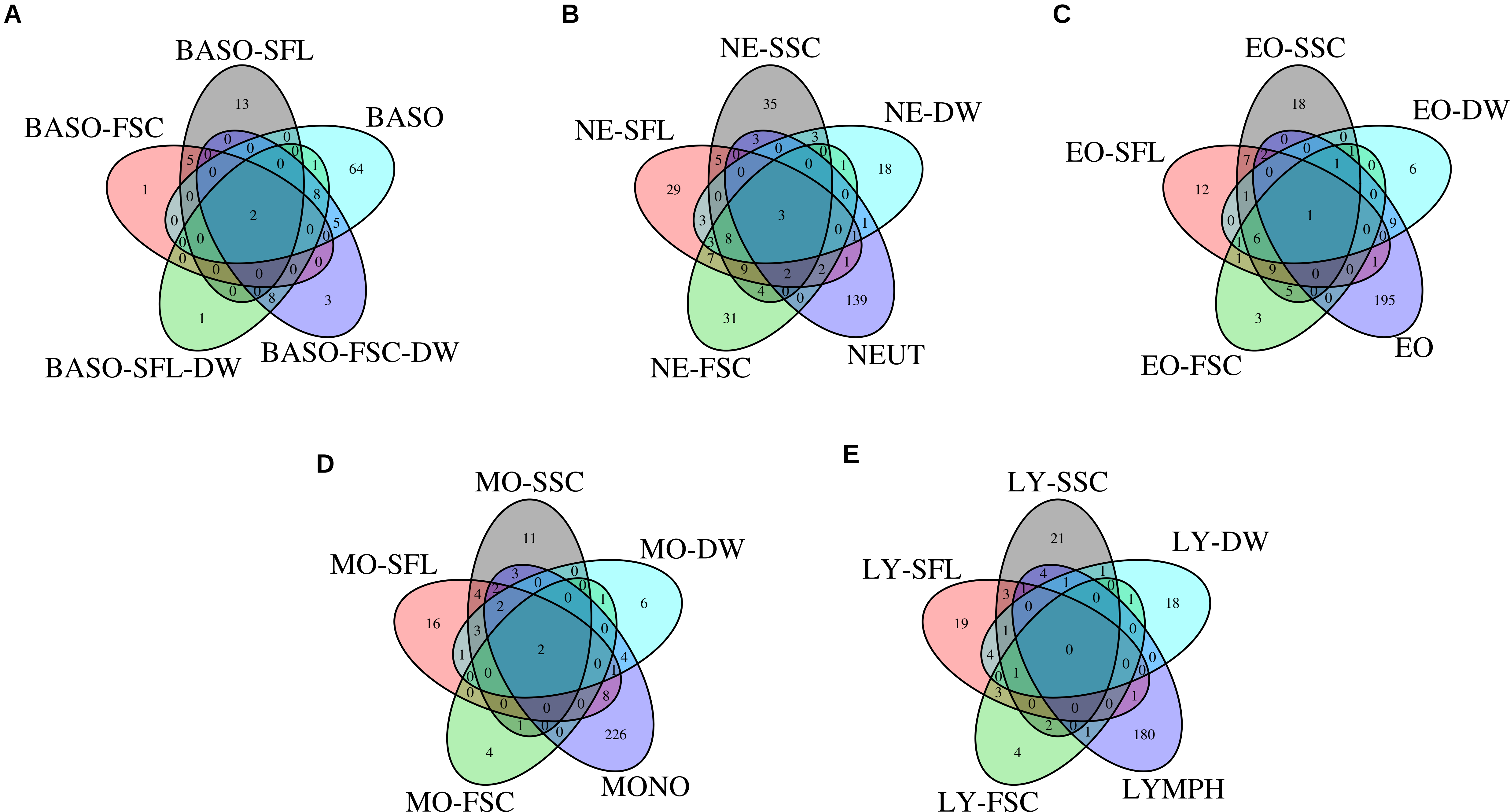
A
BASO-SFL
BASO-FSC
BASO
BASO-SFL-DW
BASO-FSC-DW
B
NE-SSC
NE-SFL
NE-DW
NE-FSC
NEUT
C
EO-SSC
EO-SFL
EO-DW
EO-FSC
EO
D
MO-SSC
MO-SFL
MO-DW
MO-FSC
MONO
E
LY-SSC
LY-SFL
LY-DW
LY-FSC
LYMPH

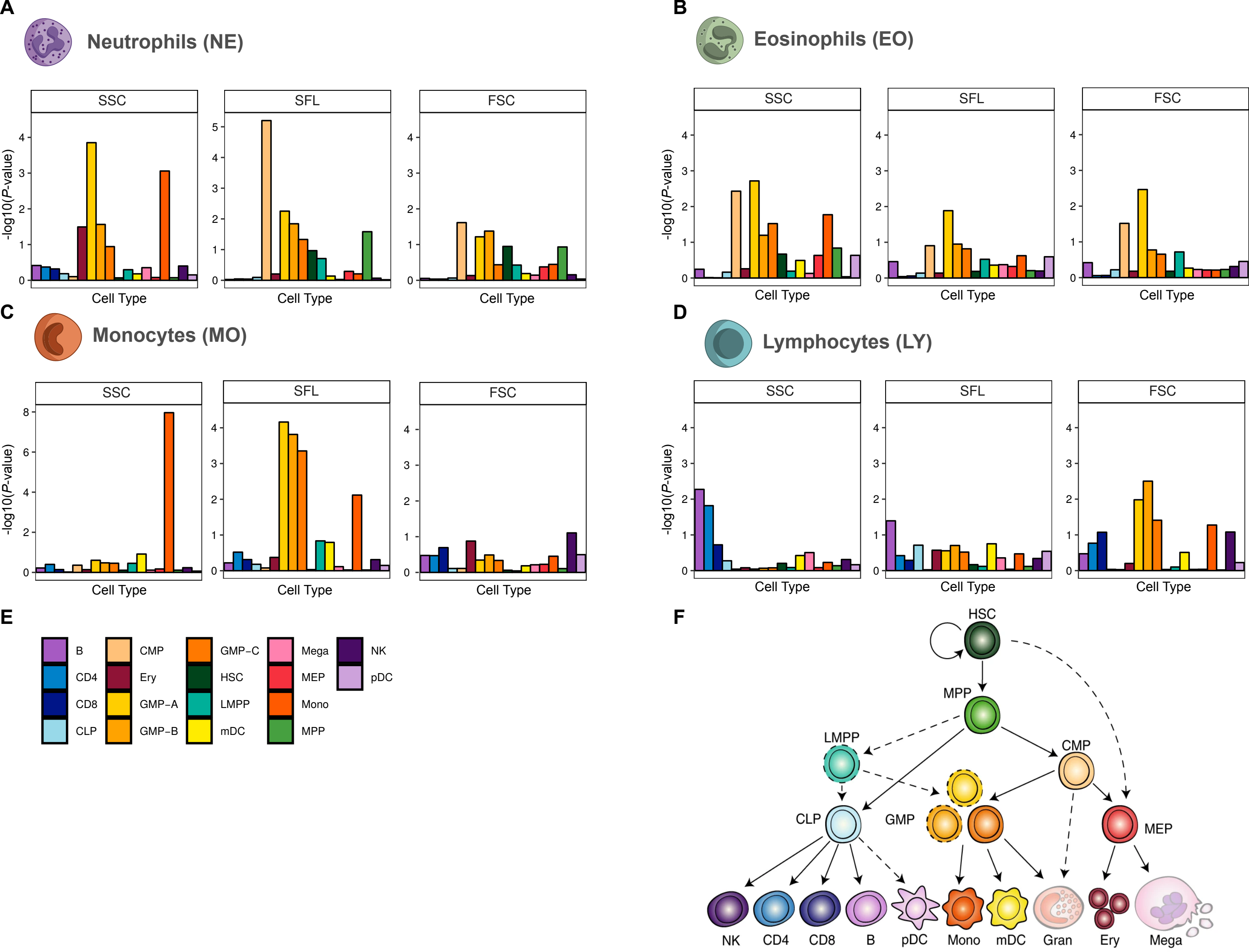
A
Neutrophils (NE)
B
Eosinophils (EO)
C
Monocytes (MO)
D
Lymphocytes (LY)
SSC
SFL
FSC
-log10(P-value)
Cell Type
E
B
CD4
CD8
CLP
CMP
Ery
GMP−A
GMP−B
GMP−C
HSC
LMPP
mDC
Mega
MEP
Mono
MPP
NK
pDC
F
HSC
MPP
LMPP
CMP
CLP
GMP
MEP
NK
CD4
CD8
B
pDC
Mono
mDC
Gran
Ery
Mega

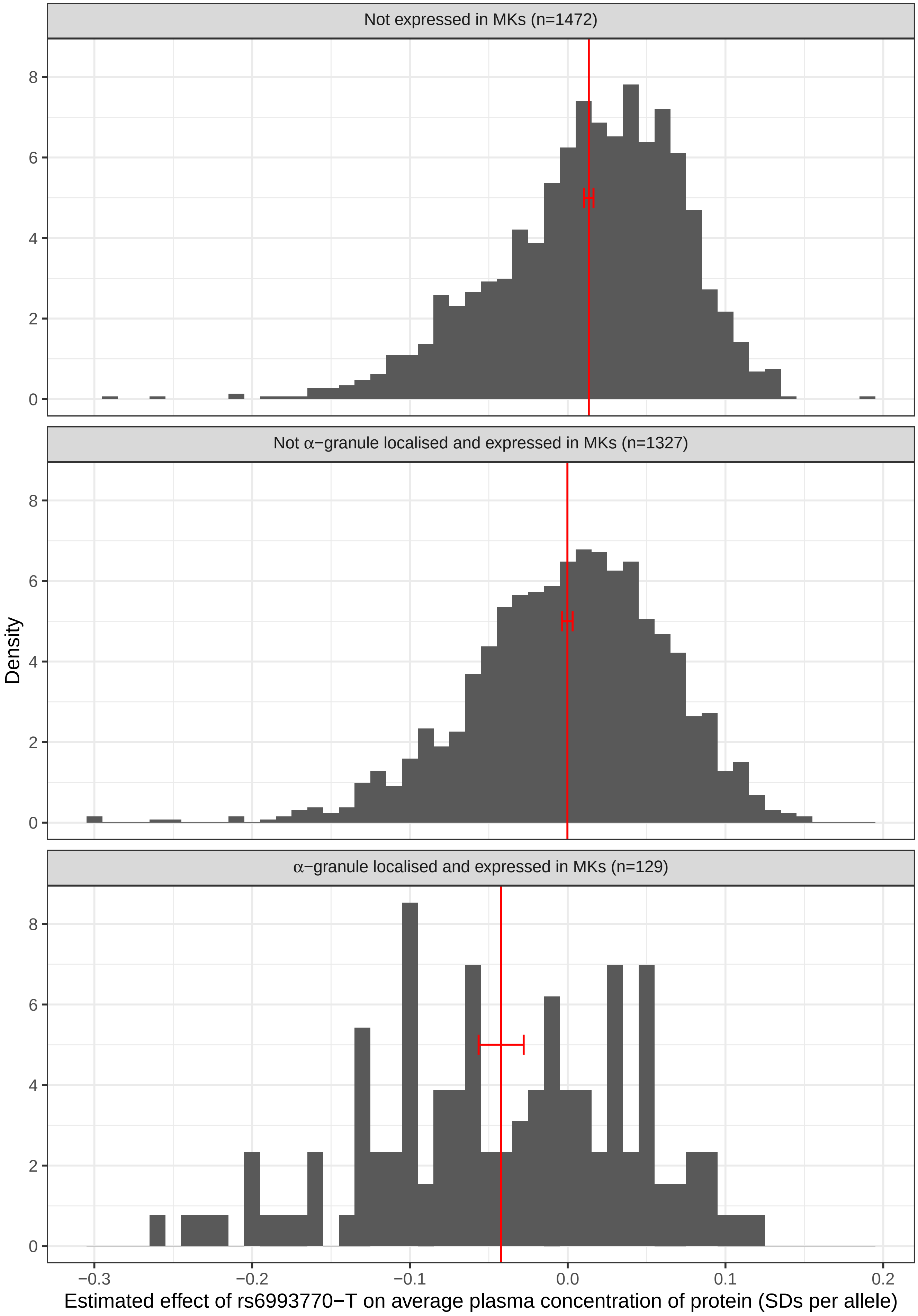

Not expressed in MKs (n=1472)
Not α−granule localised and expressed in MKs (n=1327)
α−granule localised and expressed in MKs (n=129)
Density
0
2
4
6
8
−0.3
−0.2
−0.1
0.0
0.1
0.2
Estimated effect of rs6993770−T on average plasma concentration of protein (SDs per allele)
